# Temporal complexity of fMRI is reproducible and correlates with higher order cognition

**DOI:** 10.1101/770826

**Authors:** Amir Omidvarnia, Andrew Zalesky, Sina Mansour, Dimitri Van De Ville, Graeme D. Jackson, Mangor Pedersen

## Abstract

It has been hypothesized that resting state networks (RSNs) likely display unique temporal complexity fingerprints, quantified by their multi-scale entropy patterns [1]. This is a hypothesis with a potential capacity for developing digital biomarkers of normal brain function, as well as pathological brain dysfunction. Nevertheless, a limitation of [1] was that resting state functional magnetic resonance imaging (rsfMRI) data from only 20 healthy individuals was used for the analysis. To validate this hypothesis in a larger cohort, we used rsfMRI datasets of 1000 healthy young adults from the Human Connectome Project (HCP), aged 22-35, each with four 14.4-minute rsfMRI recordings and parcellated into 379 brain regions. We quantified multi-scale entropy of rsfMRI time series averaged at different cortical and sub-cortical regions. We performed effect-size analysis on the data in 8 RSNs. Given that the morphology of multi-scale entropy is affected by the choice of its tolerance parameter (*r*) and embedding dimension (*m*), we repeated the analyses at multiple values of *r* and *m* including the values used in [1]. Our results reinforced high temporal complexity in the default mode and frontoparietal networks. Lowest temporal complexity was observed in the sub-cortical areas and limbic system. We investigated the effect of temporal resolution (determined by the repetition time *T_R_*) after downsampling of rsfMRI time series at two rates. At a low temporal resolution, we observed increased entropy and variance across datasets. Test-retest analysis showed that findings were likely reproducible across individuals over four rsfMRI runs, especially when the tolerance parameter *r* is equal to 0.5. A strong relationship was observed between temporal complexity of RSNs and fluid intelligence (people’s capacity to reason and think flexibly) through step-wise regression analysis suggesting that complex dynamics of the human brain is an important attribute of high-level brain function. Finally, the results confirmed that the relationship between functional brain connectivity strengths and rsfMRI temporal complexity changes over time scales, likely due to the regulation of neural synchrony at local and global network levels.

## 1 Introduction

The human brain is a *complex* hierarchy of modules that are dynamically interacting with each other at micro, meso and macro scales [2, 3]. Anatomically distinct, but functionally connected regions of the cortex that simultaneously fluctuate over time are referred to as *resting state networks* (*RSNs*). RSNs are intrinsic organizations of functional connectivity in the brain that are communicating with each other even in the absence of an overt cognitive tasks [4–6]. These functional brain networks can be derived from resting state functional magnetic resonance imaging (rsfMRI), and are supporting a variety of sensory, cognitive and behavioural functions [7, 8]. Perturbed functionality of RSNs contributes to a range of brain diseases including epilepsy [9], Alzheimer’s disease [10], autism [11], depression [12] and schizophrenia [13]. Although alterations of RSNs have been subject to numerous studies, characterization of their *temporal complexity* remains an open question in the brain sciences [1, 14–21]. In the context of this study, temporal complexity is referred to as a balanced dynamical behaviour between pure regularity and complete irregularity in the time domain. This is a significant challenge in modern neuroscience because temporal brain complexity may provide a quantitative view of brain function at the phenomenological level which in turn, may lead to the development of more efficient diagnostic and prognostic markers of brain diseases.

Functional co-activations associated with RSNs fluctuate over time [22, 23]. Until recently, most studies would treat functional brain connectivity as a *static* entity. The emergence of advanced neuroimaging techniques such as fast rsfMRI have opened up a new avenue for studying the dynamics of functional connectivity [24]. There is now a consensus that this dynamic behaviour resides between temporal order and disorder [25–27]. Temporal complexity of brain dynamics arises from interactions across numerous sub-components in the brain [1] and can be affected by internal and/or external factors such as sensory inputs, attention and drowsiness [28]. Revelations of this complexity include, but not limited to, self-similarity of EEG micro-state sequences [29, 30], dynamics of microscopic and mesoscopic neural networks in the brain [3, 31] and neuronal oscillations associated with different brain regions [32]. Several attempts have been made to characterize the temporal complexity of RSNs using rsfMRI data including time-frequency analysis [23], independent components analysis [33], point process analysis [34], sliding window analysis [35], phase synchrony analysis [36, 37], auto-regressive modelling [38] and nonlinear analysis [1, 18, 39, 40] (see [24] for a detailed review). An important form of temporal complexity in brain function can be observed through *multi-scale entropy* analysis of RSN dynamics [1]. Multi-scale entropy [41, 42] quantifies the rate of generation of new information in a dynamical process by computing sample entropy [43] over multiple temporal scales. Each scale provides a specific time resolution through coarse graining of the input signals. For example, random signals such as white noise have high sample entropy values at fine scales (i.e., fast fluctuations) which drop gradually in value at large scales (i.e., slow fluctuations). On the other hand, complex signals such as random walk or biosignals generate a more consistent sample entropy curve over different time scales, due to repeating information-bearing patterns across multiple time resolutions [20, 41, 42, 44].

In this paper, we investigated if the dynamics of RSNs can be differentiated based on their temporal complexity. To this end, we aimed to validate the existence of multi-scale entropy fingerprints in rsfMRI-based RSNs. This hypothesis was tested in [1] using rsfMRI datasets of 20 healthy subjects from the Human Connectome Project (HCP) [45] via multi-scale entropy analysis in four RSNs: default mode, central executive, as well as the left and right frontotemporal networks (Figure 1). Given the capacity of RSN complexity as an imaging-based marker of brain function in health and disease, we aimed to investigate this hypothesis in a larger sample cohort of 1000 rsfMRI datasets from the HCP database. We included 8 RSNs in this study, with a particular focus on to what extent rsfMRI results are dependent on the tolerance parameter *r*, embedding dimension *m* and temporal resolution of rsfMRI in multi-scale entropy analysis. We also conduced test-retest and effect-size analyses to delineate the reproducibility of RSN complexity across multiple rsfMRI scans and over subjects. We hypothesized that temporal complexity of brain function is related to higher order cognitive processes such as fluid intelligence or people’s capacity to reason and think flexibly. Lastly, we looked into the potential link between functional brain connectivity and temporal complexity of RSNs at different time scales.

**Figure 1:**
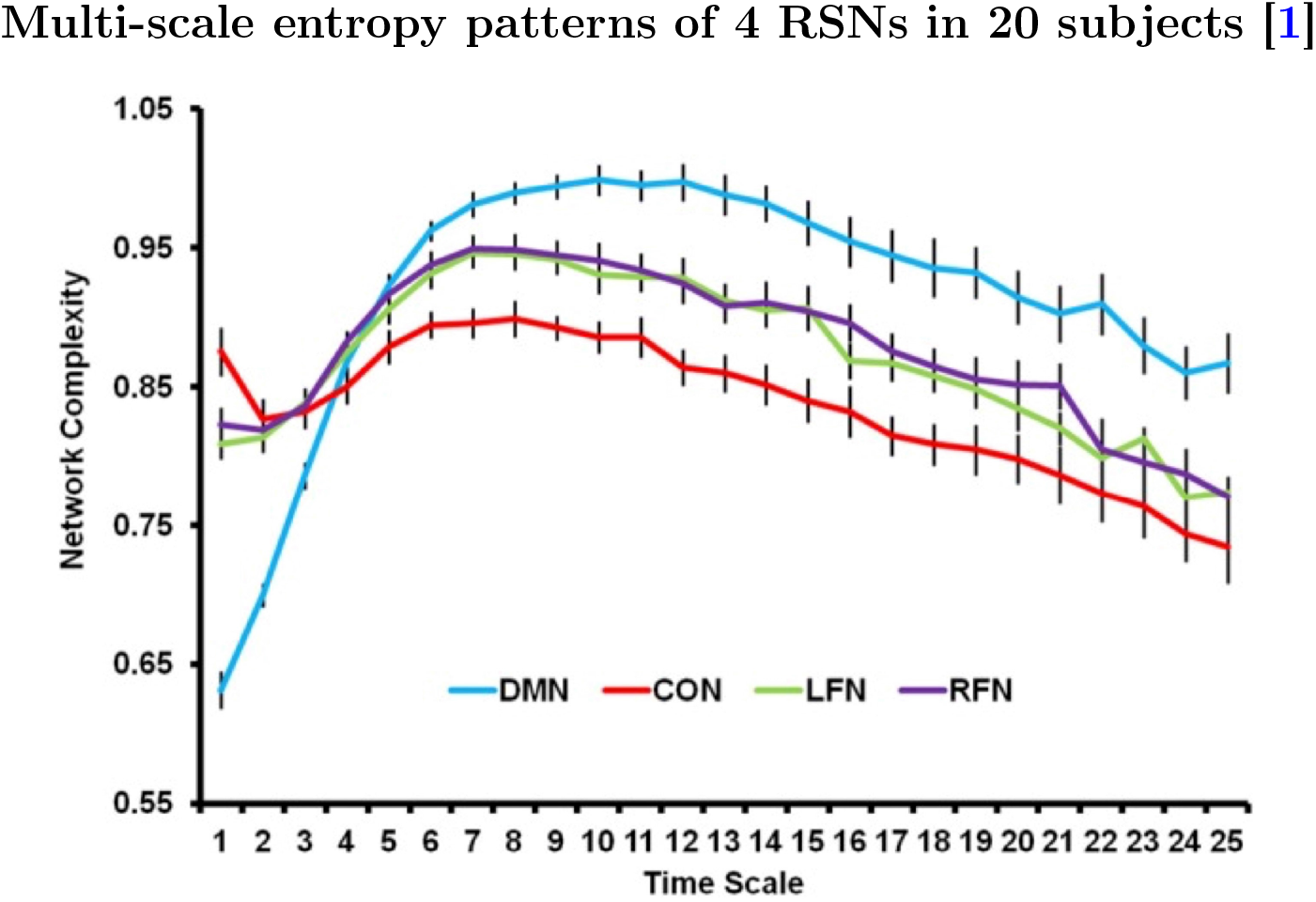
Multi-scale entropy patterns of 4 RSNs (default mode network or DMN, central executive network or CON, left frontal network or LFN, right frontal network or RFN) in 20 subjects reported in [1]. The image is taken from [1].

## 2 Materials and Methods

### 2.1 rsfMRI Data, parcellation masks and preprocessing

We used a subset of the HCP database [45] including 1000 rsfMRI datasets (N_*subj*_=1000). Each subject participated in two separate rsfMRI sessions on two different days, with two acquisitions per day, i.e., left to right and right to left slicings. We refer to these recordings as four fMRI runs throughout this paper. Each run was of length 14.4 minutes (or 1200 time points) with a voxel size of 2×2×2 millimeters the repetition time (*T_R_*) of 720 milliseconds in a 3T scanner. The following preprocessing steps were applied on each dataset: 1) echo planar imaging gradient distortion correction, 2) motion correction, 3) field bias correction, 4) spatial transformation, 5) normalization into a common Montreal Neurological Institute space [46] and 6) artefact removal using independent component analysis FIX [47]. A parcellation mask [46] was used to parcellate the gey matter into 360 cortical and 19 sub-cortical regions of interest (N_*ROI*_=379). The preprocessed datasets are publicly available at the HCP website under an Open Access Data plan agreement.

### 2.2 Multi-scale entropy analysis

While there are several definitions of signal entropy in the literature, our focus here is on multi-scale entropy analysis [41, 42]. This technique is an extended version of *sample entropy* [43] over multiple time scales.

#### 2.2.1 Sample entropy

Sample entropy [43] is a signal complexity measure which treats each short piece of an input signal **x** as a *template* to search for any *neighbouring* templates throughout the entire length of the signal. A template 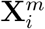 is defined as^1^:

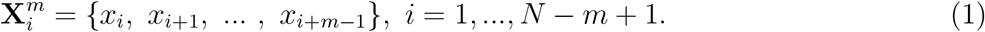

where *N* is the number of time points in **x** and *m* is the *embedding dimension* parameter. Two templates 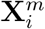 and 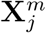 are considered as neighbours if their *Chebyshev* distance 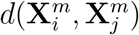 is less than a *tolerance* parameter *r*. It leads to an *r*-neighbourhood conditional probability function 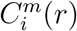 for any vector 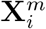 in the *m*-dimensional reconstructed phase space:

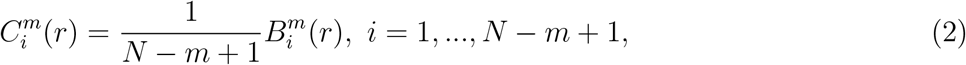

where 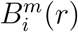 is given by:

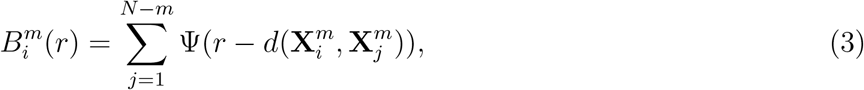

where Ψ(.) is the Heaviside function, defined as:

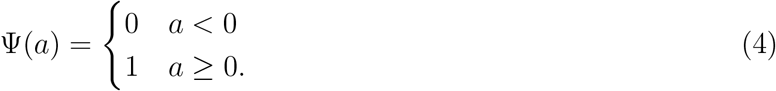

The Chebyshev distance *d* is defined as:

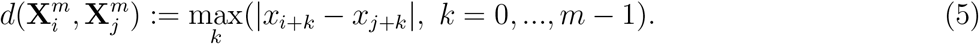

Sample entropy is then given by:

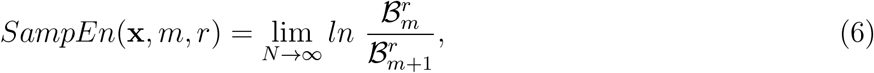

where 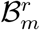 is the average of 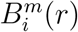 over all templates:

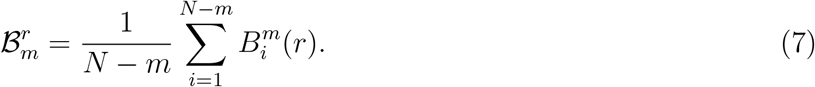

Since 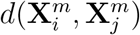 is always smaller than or equal to 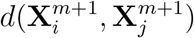, 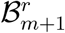 will always take smaller or equal values than 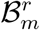. Therefore, sample entropy is always non-negative with larger values indicating less regularity [43]. The tolerance parameter *r* plays a central role in any sample entropy analysis, because it defines the probability of neighbourhood (i.e., similarity) between two templates in the reconstructed phase space. It is important to multiply *r* by the standard deviation of **x** to account for amplitude variations across different signals [18, 43]. In this study, we used the embedding dimension of *m*=2 and the tolerance parameter of *r*=0.5 for sample entropy analysis, as adapted in [1]. In addition, we used the tolerance parameter of *r*=0.15, a widely used option in the literature (see [48, 49] ans examples), as well as a range of embedding dimensions from *m*=3 to *m*=10.

#### 2.2.2 Multi-scale entropy

Multi-scale entropy extracts sample entropy after *coarse-graining* of the input signal **x** at a range of time scales *τ* [41]. A coarse-grained vector 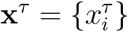 is defined as:

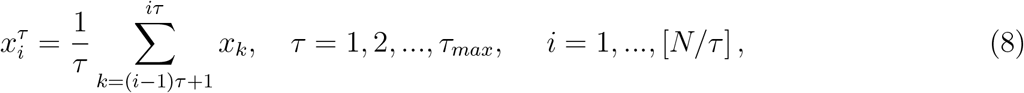

where **x**^1^ = **x**. Following [1], we set *τ_max_* to 25. At the group level, we averaged the multi-scale entropy curves over subjects and calculated the standard deviation at each scale. We also computed the *complexity index* (i.e., area under the curve) of multi-scale entropy patterns for RSNs, in all datasets.

#### 2.2.3 Complexity index

To reduce the dimensionality of multi-scale entropy patterns to a single value, a complexity index is defined as the area under each multi-scale entropy curve over all scales, divided by the maximum number of scales (i.e., *τ_max_*) [50]. For a single subject, it can be approximated with the normalized area under the multi-scale entropy curve across multiple time scales (using MATLAB’s *trapz* command):

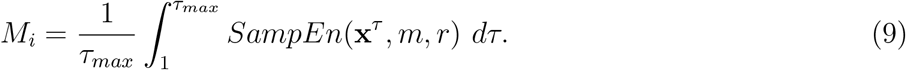

#### 2.2.4 The role of rsfMRI temporal resolution

Given the relatively short repetition time of rsfMRI time series in the HCP database (*T_R_*=0.72 seconds), we investigated to what extent the observed complex dynamics of RSNs is sensitive to rsfMRI temporal resolution. This is an important issue to check, because TR values longer than one second are common across research and clinical centres. We resembled longer *T_R_*’s in our datasets by downsampling of the rsfMRI time series in the HCP database. To this end, we calculated the complexity indices of RSNs after downsampling of the rsfMRI time series at the rates of 2 and 4, resembling the repetition times of *T_R_*=1.44 seconds and *T_R_*=2.88 seconds, respectively.

#### 2.2.5 Effect size analysis using the Hedges’ *g* measure

We quantified the difference between complex dynamics of RSNs by pair-wise effect size analysis of the complexity index distributions at three temporal resolutions (i.e., original *T_R_* of 0.72 seconds and two downsampling rates) as well as multiple combinations of the multi-scale entropy parameters (i.e., *r*=0.15, 0.5 and *m*=2 to 10). To this end, we used the Hedges’ *g_i,j_* statistic, defined as [52]:

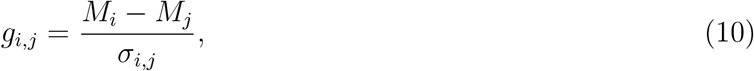

where *M_i_* and *M_j_* are the group mean complexity indices of the *i*^th^ and *j*^th^ RSNs, respectively, and *σ_i,j_* is the squared mean of the associated standard deviations computed as:

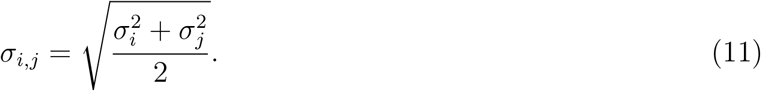

The confidence interval and *p*-value of the Hedges’ *g* measures were calculated through bootstrapping (2000 random samplings of the original time series with replacement)^2^.

#### 2.2.6 Test-retest analysis using the intra-class correlation coefficient

In order to investigate the reproducibility of multi-scale entropy patterns extracted from RSNs at different temporal resolutions, we computed *intra-class correlation* coefficient of sample entropy values at single time scales and over four rsfMRI scans (N_*run*_=4). Following [54], we chose the third intra-class correlation coefficient measure defined in [55] for test-retest analysis as:

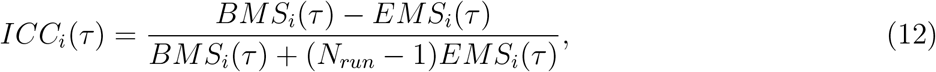

where *BMS_i_*(*τ*) and *EMS_i_*(*τ*) are the *between-subjects mean square* and the *error mean square* of sample entropy values, respectively, for the *i*^th^ RSN at the time scale *τ*. We considered the intra-class correlation coefficient values below 0.4 as poor reliability, between 0.4 and 0.6 as fair reliability and between 0.6 and 0.8 as good reliability [54].

### 2.3 Temporal complexity of RSNs and cognition

We also tested whether temporal complexity of rsfMRI is related to higher order cognition. For each subject (*N_subj_* = 1000), we selected five well-validated domain-specific behavioural variables (*N_beh_* = 5) involved in higher order cognition; *i*) the Eriksen flanker task (*Flanker_Unadj* — measuring response inhibition and task switching); *ii*) the Wisconsin Card Sorting Test (*CardSort_Unadj* — measuring cognitive flexibility); *iii*) the N-back task (*WM_Task_acc* — measuring working memory performance); *iv*) the Ravens task (*PMAT24;_A_CR* — measuring fluid intelligence); and *v*) the relational task (*Relational_Task_Acc* — measuring planning and reasoning abilities). See [56], for full information about behavioural variables included in the HCP. We defined a multiple linear regression model with *N_beh_* independent variables as follows:

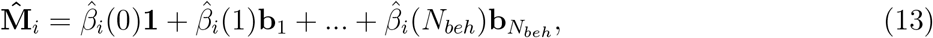

where **1** is a column vector of 1’s, 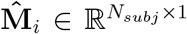 is the predicted vector of subject-specific complexity indices in the *i*^th^ RSN and 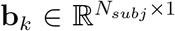 is the associated vector of *k*^th^ behavioural measures (*k* = 1,…, *N_beh_*). For each estimated coefficient 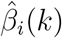, we performed a *t*-test at the significance level of 0.05 whether the coefficient is equal to zero or not. To assess whether the correlation coefficients between real complexity indices **M**_*i*_ and their predicted associates 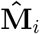 are statistically significant, we performed a permutation testing for each RSN where we permuted the order of subjects in **M**_*i*_, refitted the model and repeated this procedure for 10000 times. It led to an empirical null distribution for each network.

To assess the contribution of each behavioural variable into the temporal complexity of RSNs, we performed a bidirectional step-wise regression analysis where the independent variables were added or removed based on their importance to the fitted model in an iterative fashion at the significance level of 0.05 [57]. The procedure continues until no further improvement can be obtained in the goodness of fit of the regression model (here, at a significance level of p<0.05).

## 3 Results

### 3.1 Simulation: Multi-scale entropy analysis of color noise

To demonstrate the capacity of multi-scale entropy analysis for encoding signal dynamics, we simulated 100 realizations of four color noise signals (white, blue, pink and red) with 1200 time-points and computed their multi-scale entropy patterns (*m*=2, *r*=0.15). See Figure 2-A, B for exemplary realizations of the noise types and their associated power spectral densities. As Figure 2-C shows, multi-scale entropy curves of each noise type are distinct and can be considered as their dynamical signature. The associated complexity index values are also an informative indicator of the time-varying nature in each noise type, except for white and red noise whose complexity distributions fully overlap (Figure 2-D). Among the four, blue and white noises lead to lower complexity indices, while pink and red noises resemble complex signals due to their 1/*f^β^* spectral density functions and fractal properties [58].

**Figure 2:**
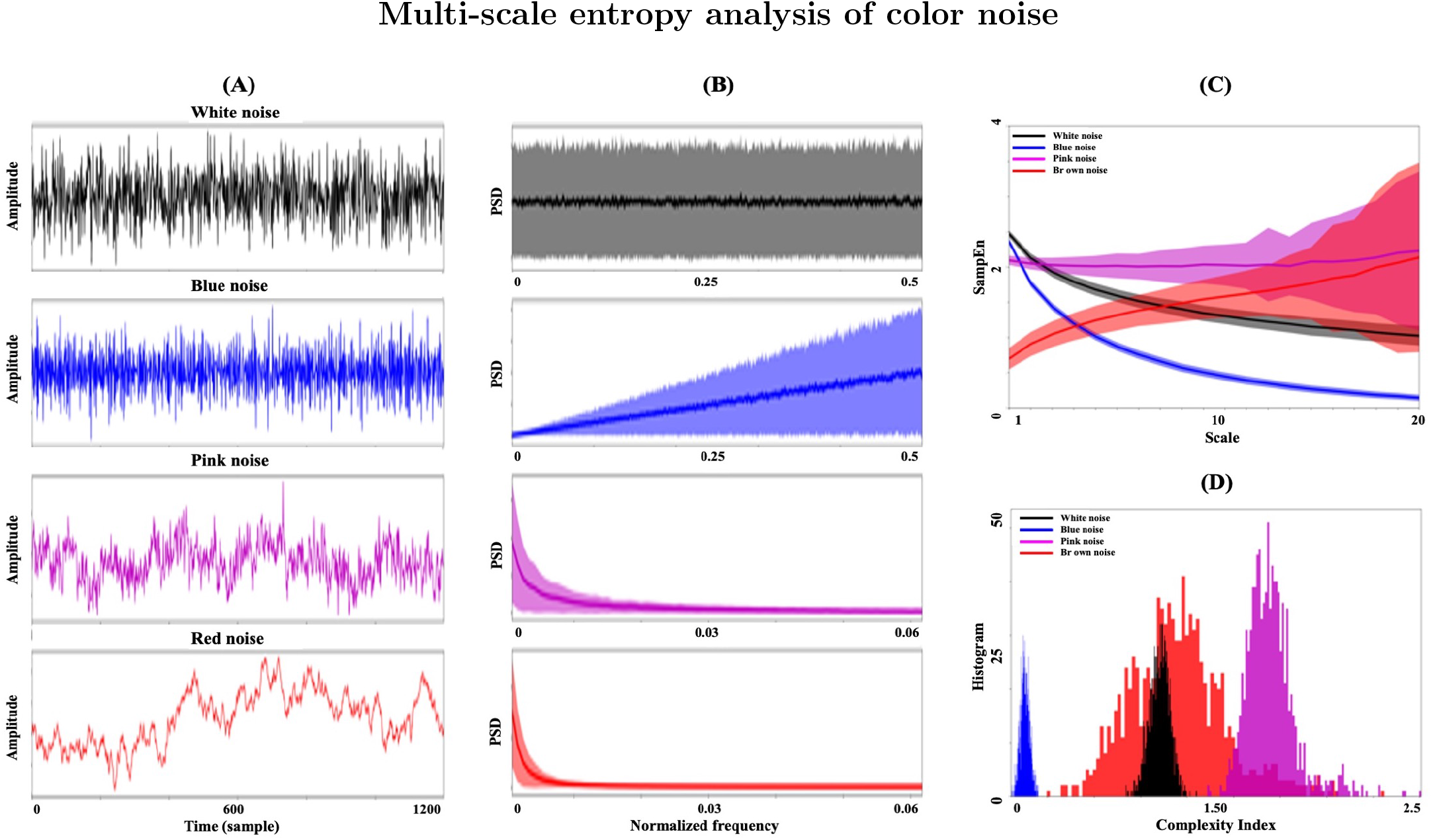
Multi-scale entropy of white noise in black color, blue noise in blue color, pink noise in pink color and red (Brown) noise in red color (*m*=2, *r*=0.15). For each noise type, 100 random realizations were generated. Column (A) Exemplary realizations in the time domain. Column (B) Shaded error bars of power spectral density functions associated with 100 realizations. (C) Distributions of multi-scale entropy patterns over 100 realizations. Shaded regions show one standard deviation from the mean curve. (D) Distributions of complexity index values.

### 3.2 RSNs are temporally complex

We observed distinct multi-scale entropy patterns between cortical and sub-cortical parts of RSNs (379 regions in total illustrated as blue and red curves in the middle rows of Figure 3 and Figure 4). A visual comparison between cortical/sub-cortical multi-scale entropy curves and simulated noise processes (Figure 2-C) suggests that the entropy patterns of higher-order RSNs are closer to the morphology of synthetic complex signals such as pink noise and red noise, while sub-cortical brain regions and limbic network are more similar to non-complex, and random, signals such as white noise and blue noise. This observation was more evident for the tolerance parameter *r*=0.5 compared to *r*=0.015 (top row of Figure 3 in contrast to the top row of Figure 4). Our multi-scale entropy analysis of RSNs at *r*=0.5, *m*=2 and *τ*=1 to 25 was in line with the findings of [1] where the default mode and frontoparietal networks were studied. Possible differences between [1] and our study may be due to the fact that we used a different brain parcellation and spatial definition of RSNs compared to *McDonough* and *Nashiro*. As the third rows of Figure 3 and Figure 4 illustrate, multi-scale entropy patterns of RSNs preserve a consistent order of complexity index across 8 RSNs with the frontoparietal (FP) and default mode networks as the most complex and the sub-cortical (SUBC) and limbic (L) networks as the least complex RSNs. Visual (VIS) and somatomotor (SM) and ventral attention (VA) networks also sit in between. This ordering remains pretty consistent after changing of the multi-scale entropy parameters, despite differences in the morphology of RSN entropy patterns. The tolerance parameter *r*=0.5 leads to a more stable morphology over different embedding dimensions *m*, while the entropy curves associated with *r*=0.15 represent considerable amount of undefined values for dimensions above 3 (*m* ≥4) and therefore, less discrimination between the temporal complexity of RSNs (bottom row in Figure 4). The undefined values of multi-scale entropy are caused by the zero values 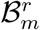 of in Eq. 6 due to the lack of neighbouring templates 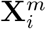 and 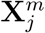 at the tolerance parameter *r*. The effect size analysis of the pair-wise comparisons across RSNs are illustrated in Figure 6 and summarized in the first columns of Table S6 (for *r*=0.5 and *m*=2) and Table S7 (for *r*=0.15 and *m*=2). According to the tables, RSNs are highly distinguishable based on their associated complexity indices at both values of *r* (Hedges’ *g* of 2.33±1.68 for *r*=0.5 and 2.55±1.72 for *r*=0.15).

**Figure 3:**
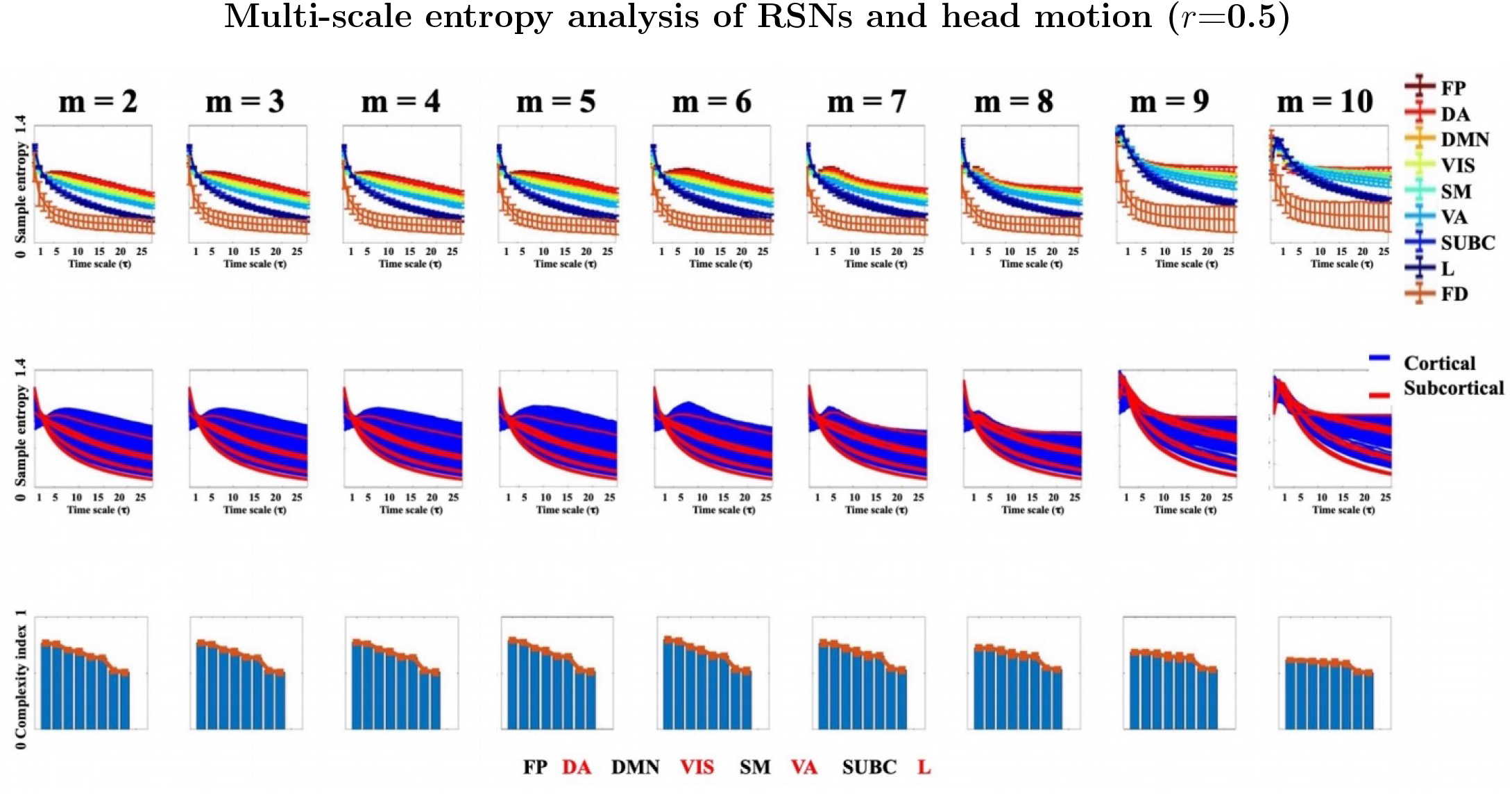
**Top row**: Multi-scale entropy patterns of 8 RSNs as well as head motion (frame-wise displacement), averaged over 1000 subjects and four rsfMRI runs, with the tolerance parameter *r*=**0.15** and a range of embedding dimensions *m* from 2 to 10. In all plots, each curve represents an average and the error bars demonstrate one standard deviation over subjects. The entropy curves have been color-coded according to their complexity indices. The multi-scale entropy curve of head motion (labeled as *FD*) has a distinct pattern compared to RSNs. **Middle row**: ROI-wise multi-scale entropy patterns, averaged over 1000 subjects and four rsfMRI runs for the same parameter sets as the first row. The entropy patterns associated with the cortical regions (*N_ROI_*=360) are in blue color and associated with the sub-cortical regions (*N_ROI_*=19 are in red color. The brain parcels were obtained according to [46]. **Bottom row**: Bar plots of the group-mean complexity indices associated with the RSN-wise multi-scale entropy patterns in the first row. The RSNs have been sorted according to their group-mean complexity indices. In all plots, the bars are labeled as follows: FP, DA, DMN, VIS, SM, VA, SUBC, L. **Abbreviation**: FP = Frontoparietal, DA = Dorsal Attention, DMN = Default mode network, VIS = Visual, SM = Sensorimotor, VA = Ventral Attention, SUBC = Sub-cortical, L = Limbic, FD = Frame-wise Displacement (extracted from head motion parameters). See [51] for the illustrations of RSNs.

**Figure 4:**
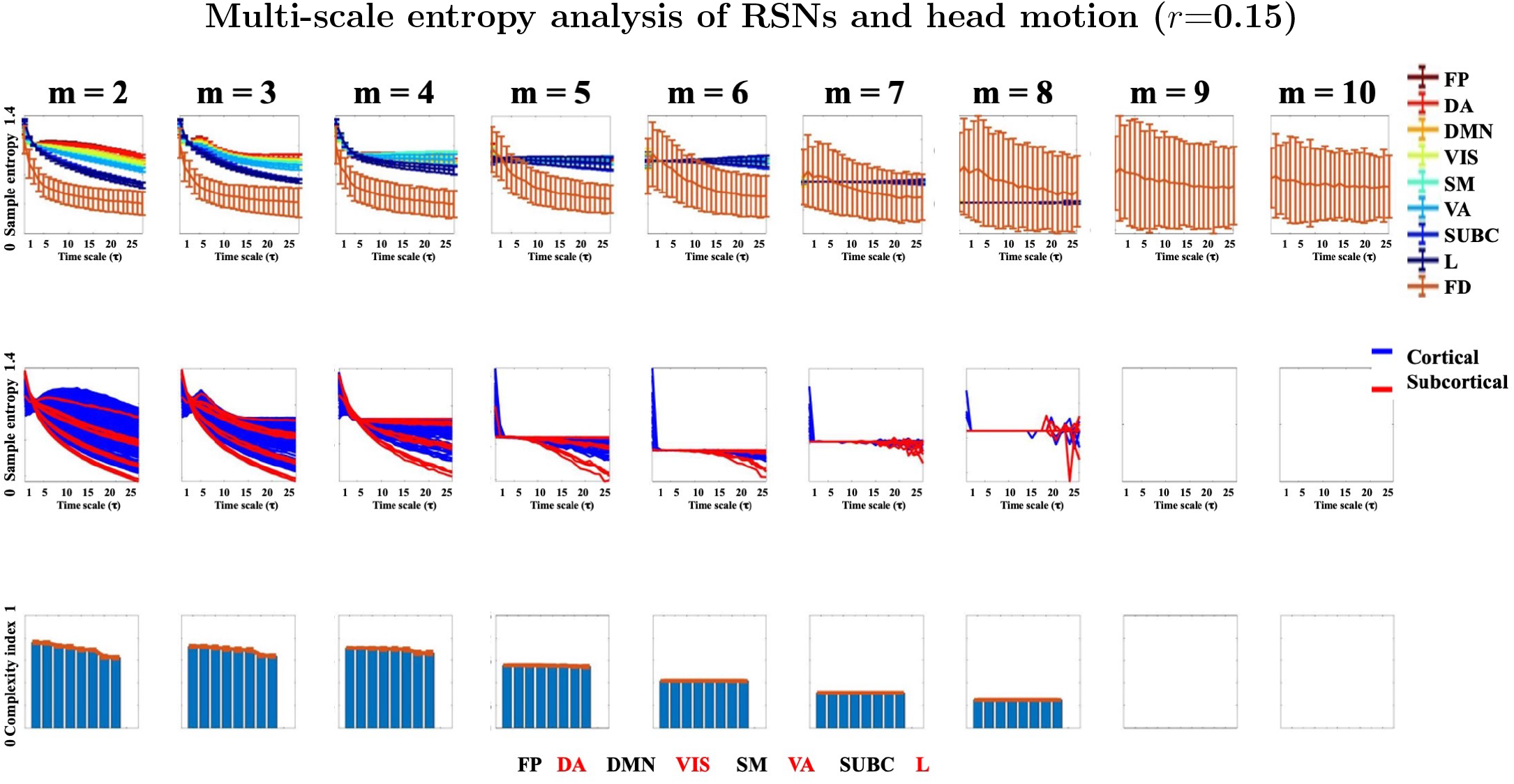
**Top row**: Multi-scale entropy patterns of 8 RSNs as well as head motion (frame-wise displacement), averaged over 1000 subjects and four rsfMRI runs, with the tolerance parameter *r*=**0.15** and a range of embedding dimensions *m* from 2 to 10. In all plots, each curve represents an average and the error bars demonstrate one standard deviation over subjects. The entropy curves have been color-coded according to their complexity indices. The multi-scale entropy curve of head motion (labeled as *FD*) has a distinct pattern compared to RSNs. **Middle row**: ROI-wise multi-scale entropy patterns, averaged over 1000 subjects and four rsfMRI runs for the same parameter sets as the first row. The entropy patterns associated with the cortical regions (*N_ROI_*=360) are in blue color and associated with the sub-cortical regions (*N_ROI_*=19 are in red color. The brain parcels were obtained according to [46]. **Bottom row**: Bar plots of the group-mean complexity indices associated with the RSN-wise multi-scale entropy patterns in the first row. The RSNs have been sorted according to their group-mean complexity indices. Blanc panels represent undefined entropy values. In all plots, the bars are labeled as follows: FP, DA, DMN, VIS, SM, VA, SUBC, L. **Abbreviation**: FP = Frontoparietal, DA = Dorsal Attention, DMN = Default mode network, VIS = Visual, SM = Sensorimotor, VA = Ventral Attention, SUBC = Sub-cortical, L = Limbic, FD = Frame-wise Displacement (extracted from head motion parameters). See [51] for the illustrations of RSNs.

### 3.3 Head motion is temporally less complex than RSN dynamics

A striking observation in the top rows of Figure 3 and Figure 4 is the distinctive multi-scale entropy pattern of head motion, quantified by frame-wise displacement of each subject during the rsfMRI runs, in contrast to the dynamics of RSNs. Frame-wise displacement of an rsfMRI recording is defined as the sum of the absolute values of the derivatives of its associated six realignment parameters [59]. As the figures suggest, head motion has a considerably lower temporal complexity than rsfMRI time series which makes it comparable with the dynamics of white and blue noises in Figure 2. This distinction is most obvious across the lower time scales (*τ* ≤10). Notably, an increase in the embedding dimension *m* has a detrimental impact on the multi-scale entropy patterns of head motion at *r*=0.15 and increases the standard deviation at each time scale drastically (see Figure 4). In fact, embedding dimensions above 3 lead to very poor outcome at *r*=0.15. From this perspective, the choice of *r*=0.5 is more appropriate for multi-scale entropy analysis of rsfMRI and head motion, as it is more robust to the changes of embedding dimension.

### 3.4 Temporal complexity of RSNs is stronger at shorter *T_R_*’s

Figure 5 to Figure S3 illustrate multi-scale entropy curves of 8 RSNs using the tolerance parameters *r*=0.5, 0.15 and embedding dimensions *m*=2,3,4, at the downsampling rates of 2 (equivalent with a *T_R_* of 1.44 seconds) and 4 (equivalent with a *T_R_* of 2.88 seconds). We observed that the morphology of entropy values was clearly influenced by the temporal resolution of the underlying data (see Figure 5 and Figure S2 versus Figure 3 for *r*=0.5 and Figure S1 and Figure S3 versus Figure 4 for *r*=0.15). This change was reflected as a decrease in the mean values of the complexity indices across RSNs (see the bottom rows in all figures). Having said that, pair-wise discrimination between the complexity index distributions of RSNs was still preserved after downsampling (see the second and third columns of Table S6 and Table S7). However, a consistent reduction was introduced to the pair-wise Hedges’ *g* statistics of effect size analysis in longer *T_R_*’s (from 2.33±1.68 to 1.83±1.33 and 1.17±0.79 for *r*=0.5 and from 2.55±1.72 to 2.06±1.32 and 0.33±0.22 for *r*=0.15). Figure 6 illustrates the color-coded Hedges’ *g* measures of rsfMRI complexity index distributions using two tolerance parameter values at three temporal resolutions and using the embedding dimension of *m*=2.

**Figure 5:**
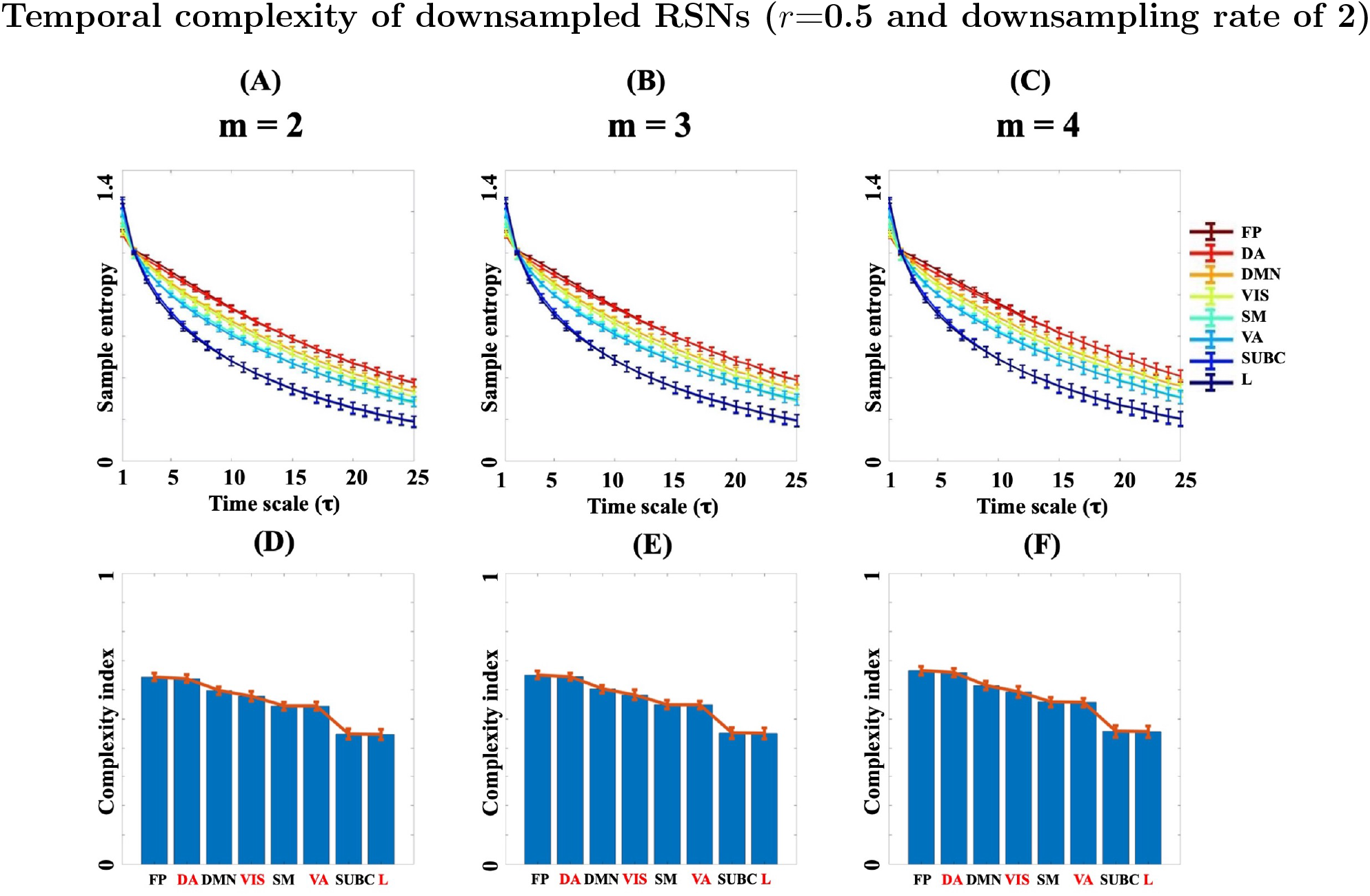
The effect of downsampling on the multi-scale entropy curves of HCP, averaged over 1000 subjects and four rsfMRI runs. (A)-(C): Error bars of multi-scale entropy curves after downsampling of rsfMRI time series **at the rate of 2 for the embedding dimensions** *m*=**2,3,4 and the tolerance parameter** *r*=**0.5**. The entropy curves have been color-coded according to their complexity indices (normalized area under their curve). (D)-(F): Mean plots of the complexity index values extracted from the multi-scale entropy curves of (A)-(C), respectively. **Abbreviation**: FP = Frontoparietal, DA = Dorsal Attention, DMN = Default mode network, VIS = Visual, SM = Sensorimotor, VA = Ventral Attention, SUBC = Sub-cortical, L = Limbic. See [51] for the illustrations of RSNs

**Figure 6:**
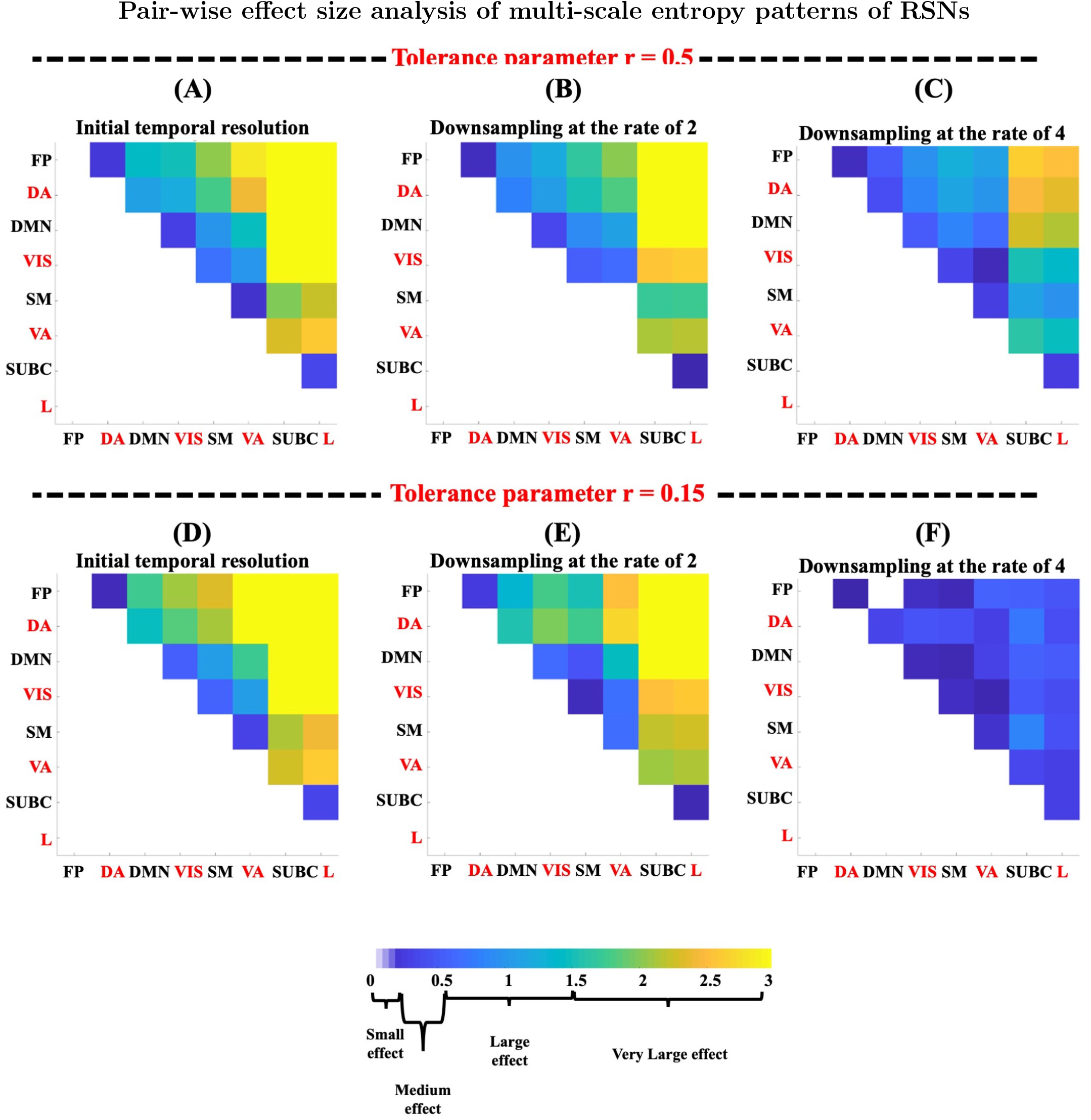
Hedges’ *g* statistics obtained from effect size analysis of the complexity index distributions calculated for each pair of RSNs. The analysis has been repeated for the embedding dimension *m*=2, the tolerance parameters (*r*=0.15,0.5) and at three downsampling scenarios (no downsampling, downsampling at the rate of 2 and downsampling at the rate of 4). The Hedges’ *g* values of less than 0.2 imply small effect, 0.2 to 0.5 are considered as medium effect, 0.5 to 1.5 are deemed as large effect and above 1.5 represent very large effect. **Abbreviation**: FP = Frontoparietal, DA = Dorsal Attention, DMN = Default mode network, VIS = Visual, SM = Sensorimotor, VA = Ventral Attention, SUBC = Sub-cortical, L = Limbic. See [51] for the illustrations of RSNs

### 3.5 Temporal complexity of RSNs is reproducible

We performed a test-retest analysis to assess whether complexity of RSNs is reproducible across different rsfMRI runs of HCP. We computed multi-scale entropy curves of 1000 datasets for four rsfMRI runs of length 14.4 minutes separately (i.e., 4×1200 *T_R_*’s). We computed the intra-class correlation coefficient of scale-dependent sample entropy values over all subjects and four sessions for 8 RSNs [51] and 25 time scales. We repeated the test-retest analysis for two tolerance parameters *r*=0.15, 0.5, three embedding dimensions *m*=2,3,4 and three rsfMRI temporal resolutions. The results are presented as color coded maps in Figure S5. As this figure shows, the tolerance parameter *r*=0.5 and *m*=2 at the original temporal resolution of rsfMRI (*T_R_*=720 msec) yielded the greatest intra-class correlation coefficient scores. At all temporal resolutions, reproducibility decreased from *r*=0.5 to *r*=0.15. Also, an increase in the embedding dimension *m* had a detrimental impact on the reproducibility of RSN complexity indices at *r*=0.15, while the values associated with *r*=0.5 were almost unchanged. Given that intraclass correlation coefficient decreases as a function of greater downsampling, it is possible that longer *T_R_*’s in the rsfMRI time series have a detrimental effect on the reproducibility of RSN complexity. Amongst the 8 RSNs, the default mode and frontoparietal networks had strongest test-retest reliability. Lowest reproducibility was seen in the sub-cortical network.

### 3.6 Temporal complexity of RSNs correlates with higher order cognition

A permutation test with 10000 randomization’s over subjects showed that correlation coefficients associated with all RSNs were above the 95^*th*^ percentile of the empirical null distributions (Figure S4). This means that the correlation between original and predicted rsfMRI complexity was statistically higher than expected by chance. We performed step-wise regression analysis between five behavioural variables and complexity indices of 8 RSNs for *m*=2, *r* = 0.15, 0.5 as well as no downsampling, downsampling at the rate of 2 and downsampling at the rate of 4 (6 different conditions in total). The results have been summarized in Figure S8 to Figure S13. As the tables show, fluid intelligence was the only winning variable in all RSNs under different scenarios. Amongst the five cognitive measures, fluid intelligence (Variable 4) displayed statistically significant (positive) regression coefficients (*β*’s) with 8 RSNs at all temporal resolutions for both tolerance parameters *r*=0.5 and *r*=0.15. To this end, we corrected each set of 8 RSN-specific *p*-values and 5 behavioural variables (i.e., 40 tests, in total) using the false discovery rate method at the significance level of *q*-value no more than 0.05. It ensures that the likelihood of false-positive results across significant regression variables stays below 5% after multiple comparisons. In other words, the association between fluid intelligence and temporal complexity of RSNs is relatively robust against the choice of tolerance parameter *r* and downsampling of the rsfMRI time series.

### 3.7 Spatial distribution of rsfMRI complexity

Figure 7 demonstrates the spatial distribution of complexity indices across 379 brain regions according to the brain parcellation of [46]. The highest complexity indices in the brain map of Figure 7-A are associated with FP, DMN and DA networks and the lowest values correspond to subcortical areas and the limbic system. More precisely, top five brain regions with the highest complexity values belong to left and right inferior parietal cortex (PGs, left and right PFm and left PF). Also, bottom five brain regions with the lowest complexity include left entorhinal cortex (EC), left and right nucleus accumbens as well as left and right pallidum. On the other hand, the lowest variability in Figure 7-B was observed across regions with the highest mean complexity including left and right inferior parietal cortex (PFm and left PGs) as well as superior parietal cortex areas associated with DMN (IP1). In contrast, the highest CI variability was associated with left inferior temporal sulcus (TE2a), left ventro-medial visual areas (VMV1), left middle temporal gyrus (TE1m), Right insular granular complex (Ig) and right lateral temporal cortex (TF). See [60] for more information about specific functions of these brain areas. This finding is consistent with our RSN specific analysis that signal complexity is highest in frontoparietal networks and DMN. Table S1 summarizes the brain regions with highest/lowest mean complexity and lowest/highest variability of complexity at the group level.

**Figure 7:**
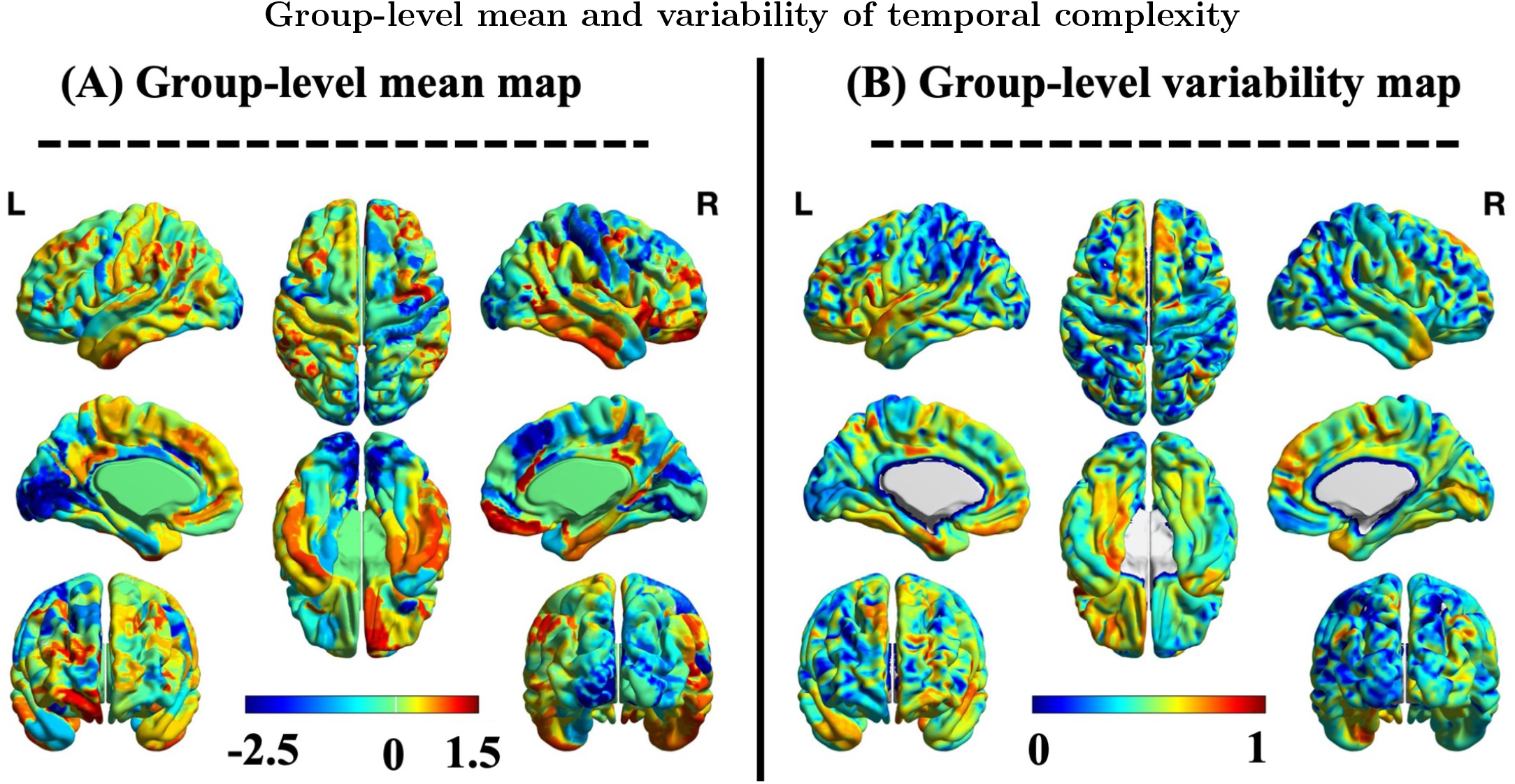
(A) Grand-mean brain map of RSN complexity indices, (B) normalized standard deviation map of RSN complexity indices. Both maps were extracted from rsfMRI datasets of 1000 HCP subjects, averaged over four resting state runs. Each dataset was parcellated using the Glasser atlas with 379 regions [60]. The complexity index is defined as the area under the curve of multi-scale entropy. The complexity indices were z-scored to aid visualization.

We also investigated the relationship between functional brain connectivity and multi-scale entropy of RSNs over time scales [1]. To this end, we extracted the *functional brain connectivity strengths* of 379 brain parcels over four rsfMRI runs leading to four brain maps for each subject. The functional connectivity strength of each ROI was defined as the sum of weights of *links* connected to that ROI. The links between ROIs were defined as the pair-wise correlation between their associated rsfMRI time series. We examined the spatial correlation between the average-run maps of sample entropy (i.e., scale-dependent multi-scale entropy) and functional connectivity strengths for each RSN separately. As Figure 8 illustrates, there is a negative correlation between functional connectivity and temporal complexity of RSNs at fine scales (i.e., between *τ*=3 for subcortical and limbic networks to *τ*=5 for frontoparietal, default mode and dorsal attention networks), while it turns to a positive correlation at coarse scales (*τ* ≥6). This observation is in line with the finding reported in [1].

**Figure 8:**
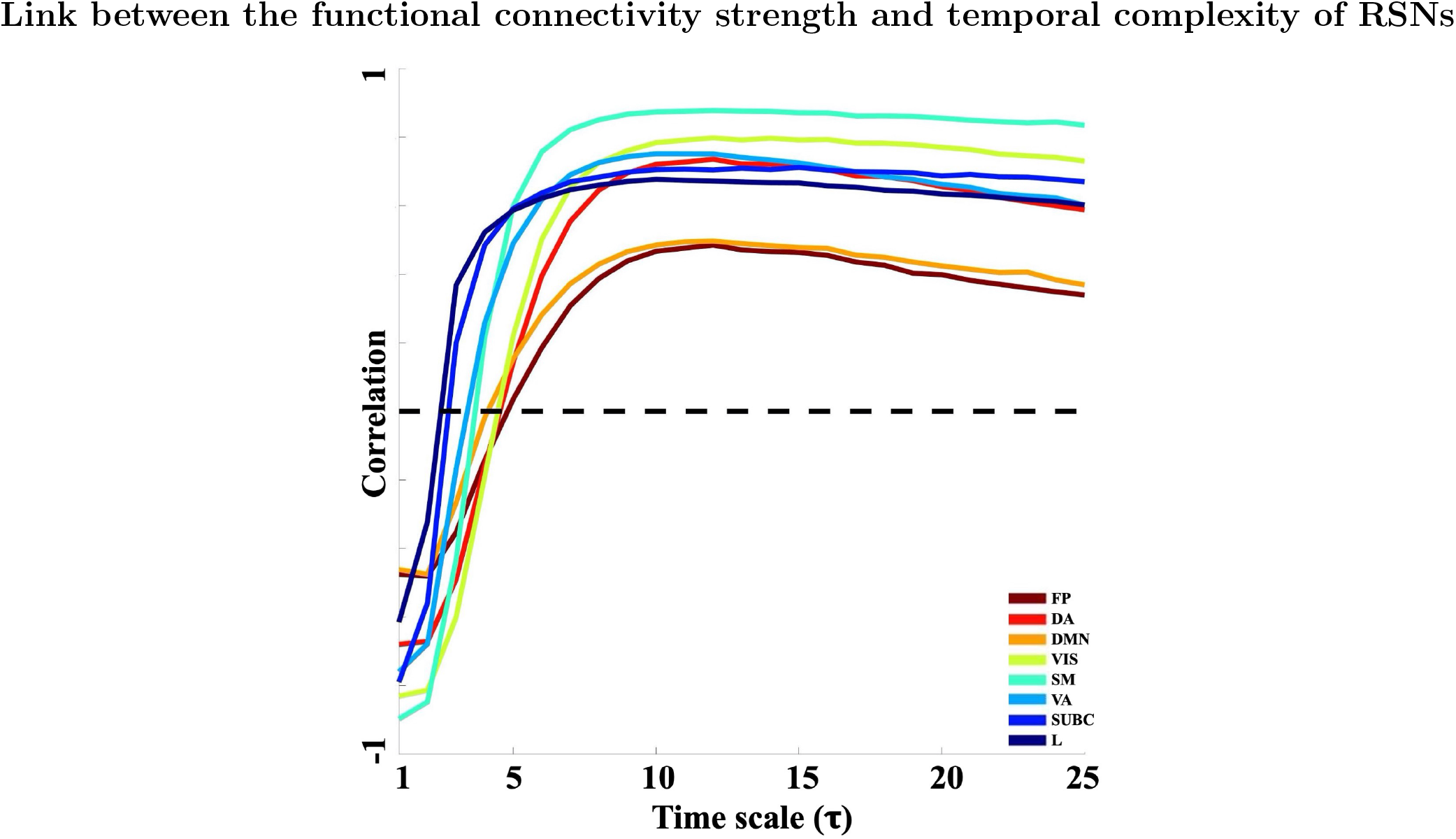
The relationship between functional connectivity strength and temporal complexity of brain regions across different RSNs. The results have been averaged over 1000 subjects and four rsfMRI runs. The networks have been colored coded according to their temporal complexity. The *x*-axis shows the time scales associated with the multi-scale entropy patterns of RSNs. The *y*-axis represents the spatial correlation between the ROI-wise functional connectivity strengths and their corresponding temporal complexity indices across different networks.

## 4 Discussion

Our study validates the hypothesis of distinct multi-scale entropy signatures in functional brain networks and reinforces the previous finding in [1]. We also build on previous research in several ways by (*i*) increasing the number of subjects from 20 to 1000, enabling a statistically more robust characterization of RSN complexity in the time domain, (*ii*) delineating temporal complexity in an additional four RSNs, (*iii*) comparing two values of the tolerance parameter r and multiple values of the embedding dimensions m for multi-scale entropy analysis, (*iv*) investigating the relationship between the temporal complexity of head motion and dynamics of RSNs, (*v*) investigating the effect of temporal resolution on the complexity of RSNs, (*vi*) analyzing the reproducibility of complex dynamics in functional networks over multiple recording sessions, and (*vii*) showing that signal complexity is related to higher-order cognitive processing.

The conceptual definition of temporal complexity may vary depending on context and data. In the context of our study, we refer to temporal complexity as a grey boundary between order and disorder over time. From this perspective, random fluctuations such as white noise have low temporal complexity, because they are completely disordered. Thus, temporal complexity is not necessarily equivalent to high unpredictability or high randomness. On the other hand, highly ordered signals such as a pure sine wave also have minimal complexity. RsfMRI sits in between these two exemplars because it represents spreading patterns of ‘structured activity’ across multiple frequency components and temporal scales that are embedded in a random background [24, 61–63]. This complex behaviour arises from functional interactions of numerous sub-components in the brain representing a balanced ‘tuning’ between order and disorder [15]. Several internal and external factors such as sensory inputs, attention, drowsiness and imagination may also ‘push’ brain dynamics towards either order or disorder, but stably [28]. It remains an open question of how to quantify ‘balanced’ fluctuations [64] in brain function and delineate their relationship with human behaviour and cognition.

The dynamics of RSNs represent a continuum of multi-scale entropy characteristics, from low complex regions across the entorhinal cortex [65] and subcortical areas including the basal ganglia have spatial patterns resembling complex noise types (i.e., pink and red noise) within frontoparietal and default mode networks [66]. See Figure 2-C in contrast to the first and second rows of Figure 3 and Figure 4. Our results suggest that the temporal complexity of RSNs is a highly discriminative feature that cannot be explained by head movement. Head motion, as quantified by frame-wise displacement, reflected a considerably lower temporal complexity than the RSN dynamics. This difference is reflected in the multi-scale entropy patterns of these signals and is significantly affected by the choice of the tolerance parameter *r* and the embedding dimension *m*. According to the results of this study, a tolerance parameter of *r* = 0.5 and an embedding dimension within the range of 2 to 5 lead to an acceptable separation between RSN dynamics and head motion as well as discrimination over RSNs. Large embedding dimensions in the multi-scale entropy analysis can increase the distance quantity 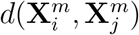 and therefore, the probability of zero outputs occurring in the Heaviside step function in Eq. 3. This is specially the case when small tolerance values are used for the multi-scale entropy analysis. Examples include the high occurrence of undefined entropy values for the combination of *r* = 0.15 and *m* ≥5 in the second and third rows of Figure 4. As presented in Tables S8 to S13, fluid intelligence seems to have the strongest linear relationship with the temporal complexity of rsfMRI. This behavioural measures refers to people’s ability to provide logical solutions to specific problems, in novel situations where acquired knowledge cannot be retrieved [67].

Vigilance is another important aspect of cognition whose influence on RSNs has been studied before [68–70]. It has been hypothesized that the temporal behaviour of RSNs is influenced by variations in vigilance [68]. Maintaining a constant level of wakefulness is difficult during resting state experiments, although HCP subjects are instructed to keep awake and visually fixate on a cross on a screen. However, it is still important to consider the potential impact of vigilance fluctuations on the multiscale entropy patterns of RSNs and their associated complexly indices. This can also influence the interpretations of which functional brain networks have more reliable brain complexity dynamics. The relationship between vigilance and temporal complexity of RSNs is regarded to future work.

Multi-scale entropy patterns provide a more comprehensive picture about brain complexity than sample entropy at single time scales. In fact, single-scale sample entropy analysis could lead to misleading interpretations about the complexity of brain regions and functional networks, as demonstrated in the top rows of Figure 3 and Figure 4 the entropy values associated with different RSNs may get reverse over large scales (for example, before and after *τ*=3 at the tolerance parameter *r*=0.5 and the embedding dimension *m*=2). The distinction in complexity between cortex and subcortex is likely related to lower temporal signal to noise ratio in rsfMRI time series within subcortical nuclei. This may be due to a higher vulnerability to thermal noise related to MRI system electronics, gradient switching artifact and physiological noise including cardiac pulsations and respiratory activity [71]. This can be further investigated using 7T data, or multi-echo data, by testing whether this distinction remains in data where the sub-cortical signal to noise ratio is improved. Figure 3 and Figure 4 show that multi-scale entropy curves, and their associated complexity indices, are considerably affected by the choice of the tolerance parameter *r* and embedding dimension *m* (see [72] for another in-depth investigation of the role of *r* and *m* in sample entropy). As the bottom rows of the figures suggest, the *relative network ordering* of complexity indices is a more consistent discriminative feature across RSNs. The effect size of complex signatures across RSNs decreases at smaller *r*’s (note the difference between the upper and lower rows of Figure 6) and at lower temporal resolutions (note the systematic reduction from left to right in Figure 6). Having said that, almost all of pair-wise Hedges’ *g* statistics remain statistically significant after bootstrapping (see Table S6 and Table S7), also due to our large sample cohort. The embedding dimension *m* is an influential factor in multi-scale entropy analysis which controls the dimensionality of the reconstructed phase space [72]. For a given embedding dimension *m*, a tolerance *r*, and a single time scale *τ*, multi-scale entropy estimates the average logarithm of the probability that if two segments of length *m* in the data have distance *r* then two segments of length *m* +1 also have distance *r*. We tested a range of values for *m* throughout the study and found a consistent estimate of multi-scale entropy for the tolerance parameter *r*=0.5 up to *m* = 10 and a severe loss of the estimates for *r*=0.15 for *m* ≥4 at the original temporal resolution of rsfMRI, i.e., *T_R_*=720 msec (Figure 3 vs. Figure 4). The mean RSN complexity indices in the lower rows of Figure 5 as well as Figure S1 to Figure S3 suggest that at longer *T_R_*’s the embedding dimension above *m*=3 may reduce the separability of RSNs (in particular, as the tolerance parameter *r*=0.15). Since HCP datasets consist of four rsfMRI recording sessions per subject, we were in a good position to perform a test-retest analysis of network-specific multi-scale entropy. Figure S5 illustrates the finding in terms of two color coded maps based on the intra-class correlation coefficient, a measure of reproducibility, extracted from network-specific sample entropy distributions at single time scales. As the figure suggests, sample entropy values over fine time scales (*τ* ≤5) are more repeatable than the values extracted at large scales. This finding was not surprising because coarse-graining step of the multi-scale entropy analysis at large *τ* can remove original information from rsfMRI time series and reduce them into a series of random fluctuations.

The biological underpinnings of multi-scale entropy has been subject to several studies in the recent years (e.g., [1, 21, 73]). Ghanbari *et al*. [74] hypothesized that more predictable neural signals establish synchronized links between remote brain regions and, therefore, facilitate long-range information processing of functional brain networks. Also, increasingly random signals are related to the local firing of neural populations. This dichotomy has been also reported for coarse-fine time scales of multi-scale entropy: fine scale values correspond to local information processing of brain networks, while coarse scale values deal with long-range communications [1, 21, 40, 75]. Functional brain connectivity may play an important role here. In fact, functional brain connectivity and temporal complexity of RSNs represent a scale-dependant relationship with a negative correlation at fine scales (small values of *τ*) and a positive correlation at coarse scales (large values of *τ*) [1]. Our results not only reinforce this finding (see Figure 8), but also they suggest that the brain regions with highest mean temporal complexity are mainly located across the default mode network, frontoparietal network and dorsal attention network (see Figure 7). These regions have also been reported as having high *participation coefficients* in the functional brain networks and therefore, playing as *connector nodes* in the brain [76, 77]. The overlap between the participation coefficient and temporal complexity brain maps may suggest a link between RSN complexity and integration of various cognitive functions in the brain. As Figure 7 illustrates, the temporal complexity patterns of rsfMRI are asymmetrical across the brain (e.g. frontal, temporal, and primary motor regions). This observation is in line with the lateralization of human brain organization and cognition [78].

The impact of rsfMRI preprocessing has been subject to extensive research (see [79–82], for examples). However, there is still no complete agreement on the most appropriate’ rsfMRI preprocessing pipeline, as this depends on several factors such as the MRI scanner type, scanning parameters, subject-specific movement artifacts, health conditions and nature of the study (e.g., whether it is a resting state study or an event-related study). The rsfMRI datasets of this study were preprocessed using a customized pipeline for the HCP project that includes ICA-FIX [46]. There is evidence suggesting that ICA-FIX is robust in reducing artefacts in large rsfMRI datasets. That is why it has been the recommended rsfMRI preprocessing pipeline by the HCP because it allows for ‘combined cortical surface and subcortical volume analysis’ [46], a requirement for the cortical-subcortical multi-scale entropy analysis of this study. Also, motion artefact removal using the FIX-ICA method [47, 83] has been shown to result in significantly improved RSN reproducibility, regardless of the recording conditions [84]. The choice of brain parcellation is another impactful factor in the temporal complexity analysis of rsfMRI which defines the spatial extent and morphology of ROIs and RSNs. It is important to use non-overlapping brain parcels (e.g., the brain atlas [60] used in tis study) RSNs in order to avoid any interference of complex dynamics across brain regions. An example of an overlapping brain parcellation is the definition of ROIs based on principal components of brain function.

A limitation of multi-scale entropy for temporal complexity analysis of functional brain networks originates from the general framework of sample and multi-scale entropy analyses as uni-variate methods. This means that the input signal to these measures is always one-dimensional. In the context of this study, ROI-specific multi-scale entropy patterns only capture the local aspects of regionally averaged rsfMRI signals and do not measure interactions between brain regions necessarily. Although we considered pair-wise relationships between the complexity distributions of RSNs (see Figures S6 and S7), it is not a substitute for multivariate complexity measures. It also speaks to the necessity of developing multivariate versions of sample/multi-scale entropy measures which can deal with a more global picture of dynamic brain function at once (for example, see [85]).

## 5 Conclusion

Functional brain networks represent distinctive signatures of temporal complexity which can be quantified through multi-scale entropy analysis of rsfMRI. This observation is robust over a large cohort of healthy subjects and reproducible over rsfMRI recording sessions. Head motion has a significantly lower temporal complexity than RSNs. Also, there is likely a strong relationship between temporal complexity of RSNs and higher-order cognition (fluid intelligence).

## Acknowledgement

AO acknowledges financial support through the Eurotech Postdoc Program, co-funded by the European Commission under its framework program Horizon 2020 (Grant Agreement number 754462). This study was supported by the National Health and Medical Research Council (NHMRC) of Australia (no 628952). The Florey Institute of Neuroscience and Mental Health acknowledges the strong support from the Victorian Government and in particular the funding from the Operational Infrastructure Support Grant. We also acknowledge the facilities, and the scientific and technical assistance of the National Imaging Facility (NIF) at the Florey node and The Victorian Biomedical Imaging Capability (VBIC). GJ is supported by an NHMRC practitioner’s fellowship (no 1060312). The primary rsfMRI data in this study was provided by the Human Connectome Project, WUMinn Consortium (1U54MH091657; Principal Investigators: David Van Essen and Kamil Ugurbil) funded by the 16 National Institutes of Health (NIH) institutes and centers that support the NIH Blueprint for Neuroscience Research; and by the McDonnell Center for Systems Neuroscience at Washington University.

## 6 Supplementary materials

**Figure S1:**
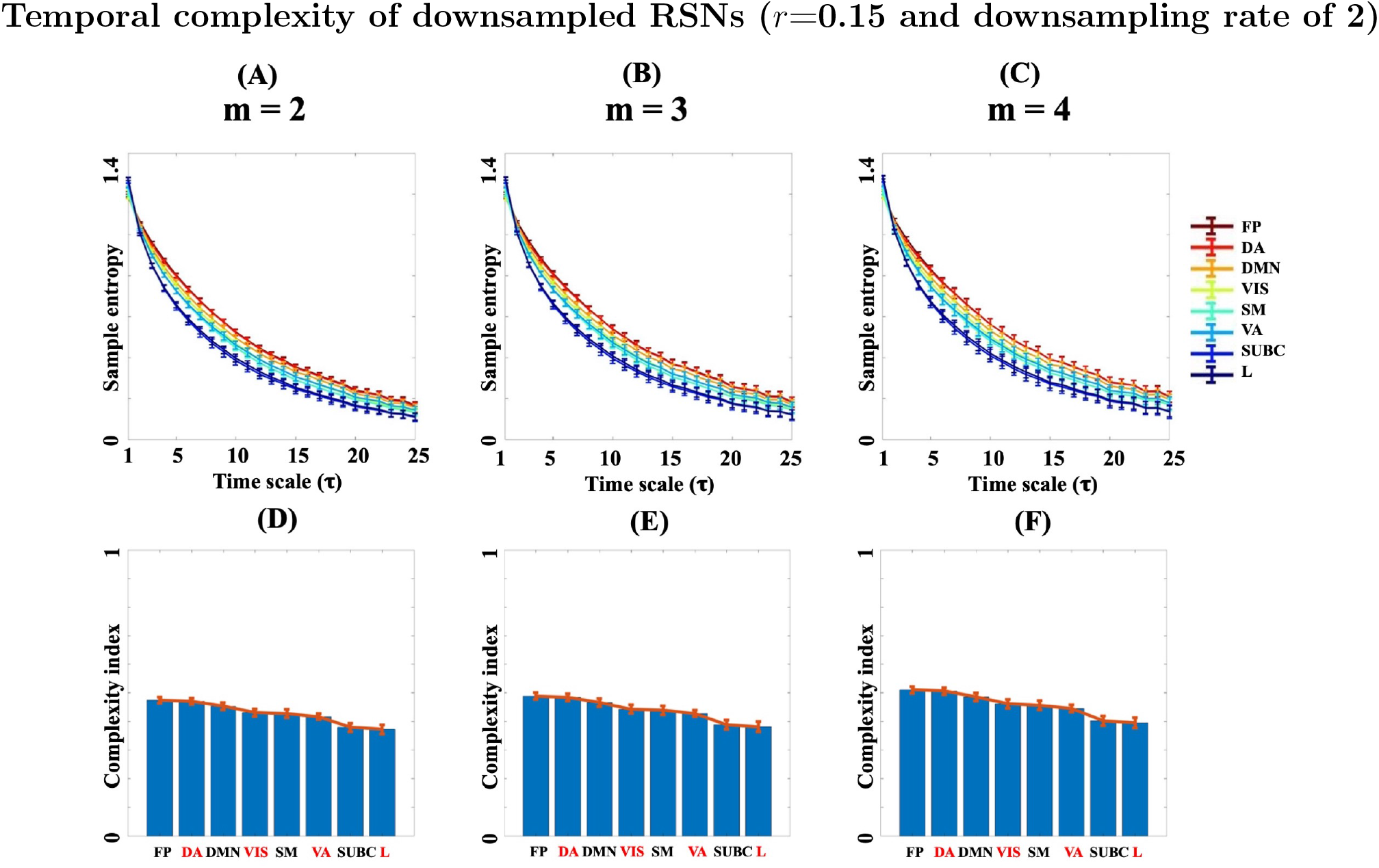
The effect of downsampling on the multi-scale entropy curves of HCP, averaged over 1000 subjects and four rsfMRI runs. (A)-(C): Error bars of multi-scale entropy curves after downsampling of rsfMRI time series **at the rate of 2 for the embedding dimensions** *m*=**2,3,4 and the tolerance parameter** *r*=**0.15**. The entropy curves have been color-coded according to their complexity indices (normalized area under their curve). (D)-(F): Mean plots of the complexity index values extracted from the multi-scale entropy curves of (A)-(C), respectively. **Abbreviation**: FP = Frontoparietal, DA = Dorsal Attention, DMN = Default mode network, VIS = Visual, SM = Sensorimotor, VA = Ventral Attention, SUBC = Sub-cortical, L = Limbic. See [51] for the illustrations of RSNs

**Figure S2:**
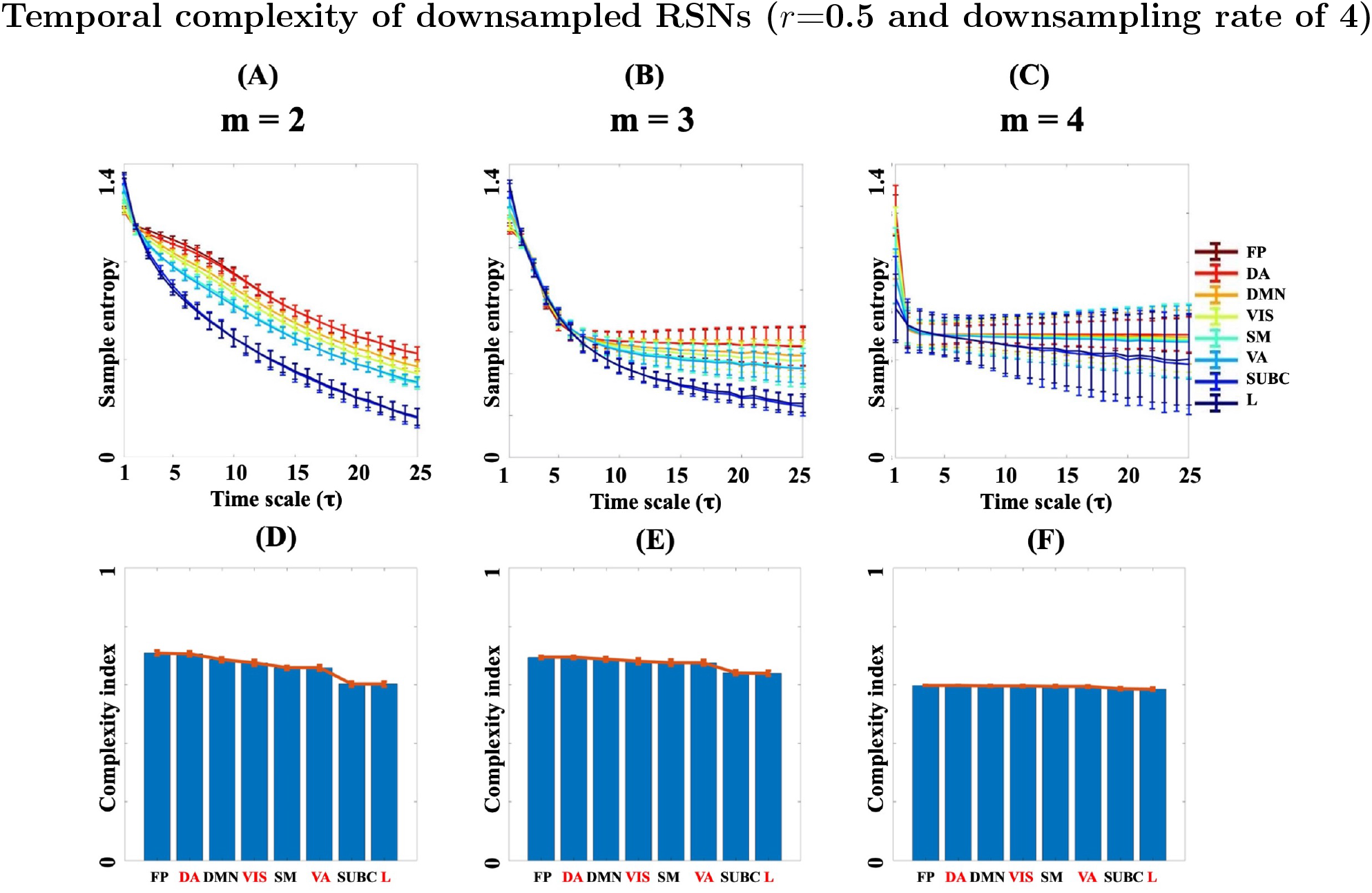
The effect of downsampling on the multi-scale entropy curves of HCP, averaged over 1000 subjects and four rsfMRI runs. (A)-(C): Error bars of multi-scale entropy curves after downsampling of rsfMRI time series **at the rate of 4 for the embedding dimensions** *m*=**2,3,4 and the tolerance parameter** *r*=**0.5**. The entropy curves have been color-coded according to their complexity indices (normalized area under their curve). (D)-(F): Mean plots of the complexity index values extracted from the multi-scale entropy curves of (A)-(C), respectively. **Abbreviation**: FP = Frontoparietal, DA = Dorsal Attention, DMN = Default mode network, VIS = Visual, SM = Sensorimotor, VA = Ventral Attention, SUBC = Sub-cortical, L = Limbic. See [51] for the illustrations of RSNs

**Table S1:** Brain regions with highest and lowest mean/variability temporal complexity indices at the group-level. The regions are defined according to the Glasser atlas [60]. In the table, group-averaged *complexity index* (i.e., the normalized area under the multi-scale entropy patterns) is denoted as *m_CI_* and normalized variability of *complexity indices* across subjects is denoted as *σ_CI_*. **Abbreviation**: EC = entorhinal cortex, PGs = in the inferior parietal cortex, PFm = in the inferior parietal cortex, PF = in the inferior parietal cortex, TE2a = inferior temporal sulcus, VMV1 = ventro-medial visual areas, TE1m = middle temporal gyrus, Ig = insular granular complex, TF = lateral temporal cortex.

| Group-mean CI |  |  | Variability of CI |  |  |
| --- | --- | --- | --- | --- | --- |
| Rank | Highest $m_{CI}$ | Lowest $m_{CI}$ | Rank | Highest $\sigma_{CI}$ | Lowest $\sigma_{CI}$ |
| 1 | Left PGs | Left EC | 1 | Left TE2a | Right PF |
| 2 | Right PFm | Right Accumbens | 2 | Left VMV1 | Right IP1 |
| 3 | Left PFm | Left Accumbens | 3 | Left TE1m | Left PFm |
| 4 | Right PGs | Right Pallidum | 4 | Right Ig | Left PGs |
| 5 | Left PF | Left Pallidum | 5 | Right TF | Right PFm |

**Figure S3:**
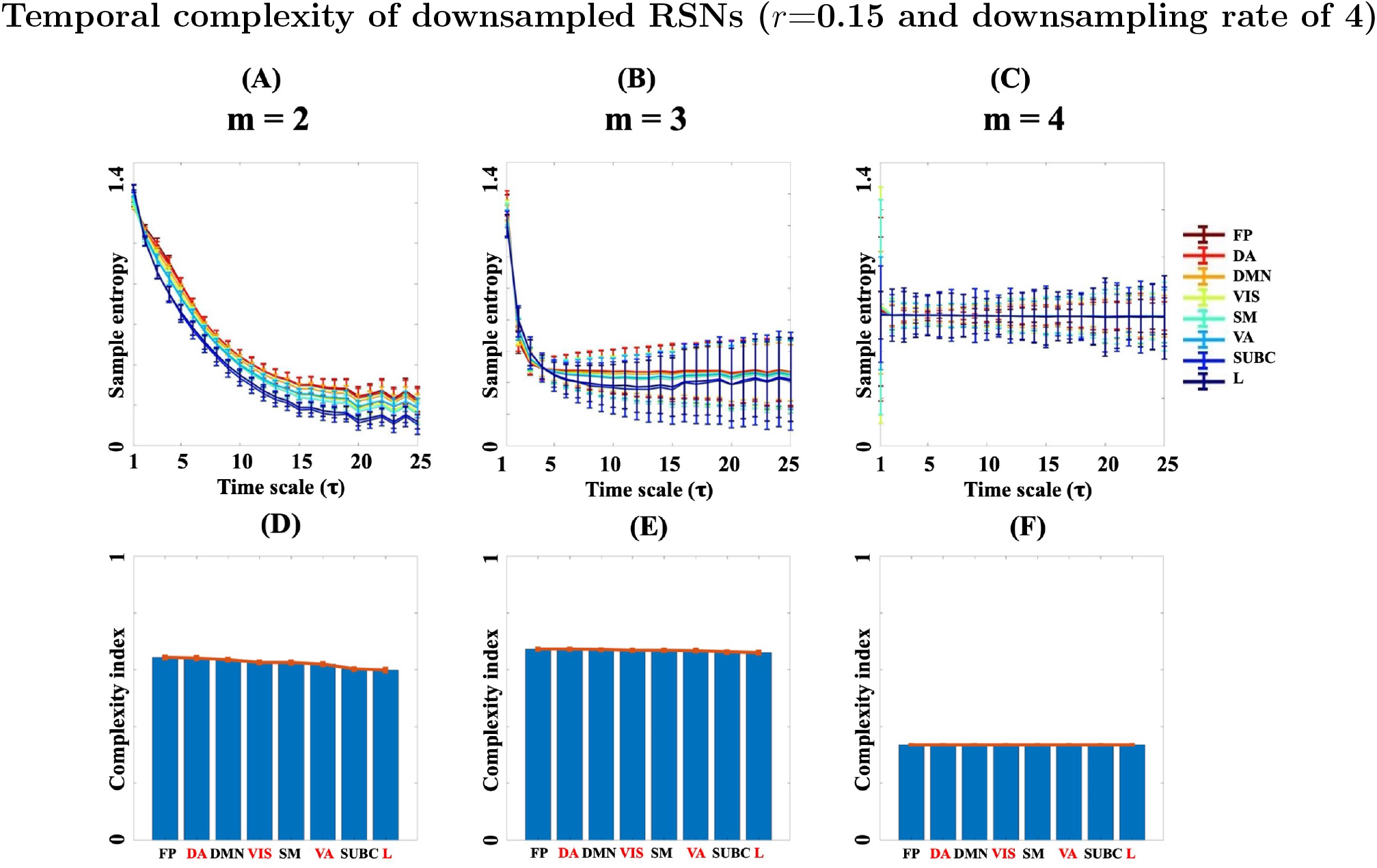
The effect of downsampling on the multi-scale entropy curves of HCP, averaged over 1000 subjects and four rsfMRI runs. (A)-(C): Error bars of multi-scale entropy curves after downsampling of rsfMRI time series **at the rate of 4 for the embedding dimensions** *m*=**2,3,4 and the tolerance parameter** *r*=**0.15**. The entropy curves have been color-coded according to their complexity indices (normalized area under their curve). (D)-(F): Mean plots of the complexity index values extracted from the multi-scale entropy curves of (A)-(C), respectively. **Abbreviation**: FP = Frontoparietal, DA = Dorsal Attention, DMN = Default mode network, VIS = Visual, SM = Sensorimotor, VA = Ventral Attention, SUBC = Sub-cortical, L = Limbic. See [51] for the illustrations of RSNs

**Figure S4:**
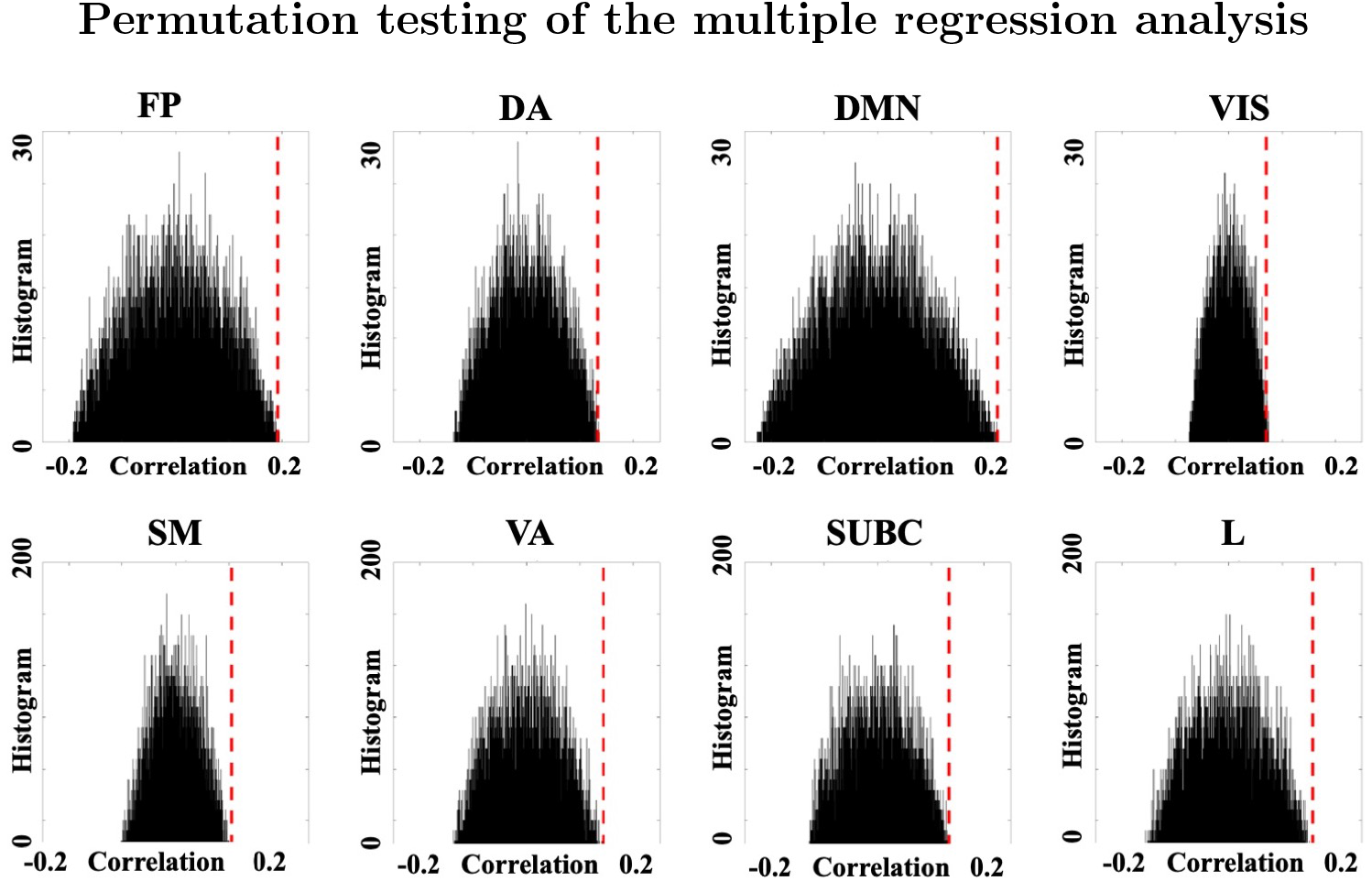
Empirical null distributions of the Spearman correlation coefficients obtained through a permutation testing with 10000 shuffling over subjects through multiple regression analysis between temporal complexity of RSNs (as output of the model) and five behavioural variables (as predictors). The dashed vertical line in each panel illustrates the Spearman correlation coefficient between the original complexity values (without shuffling) and their predictions. **Abbreviation**: FP = Frontoparietal, DA = Dorsal Attention, DMN = Default mode network, VIS = Visual, SM = Sensorimotor, VA = Ventral Attention, SUBC = Sub-cortical, L = Limbic. See [51] for the illustrations of RSNs.

**Figure S5:**
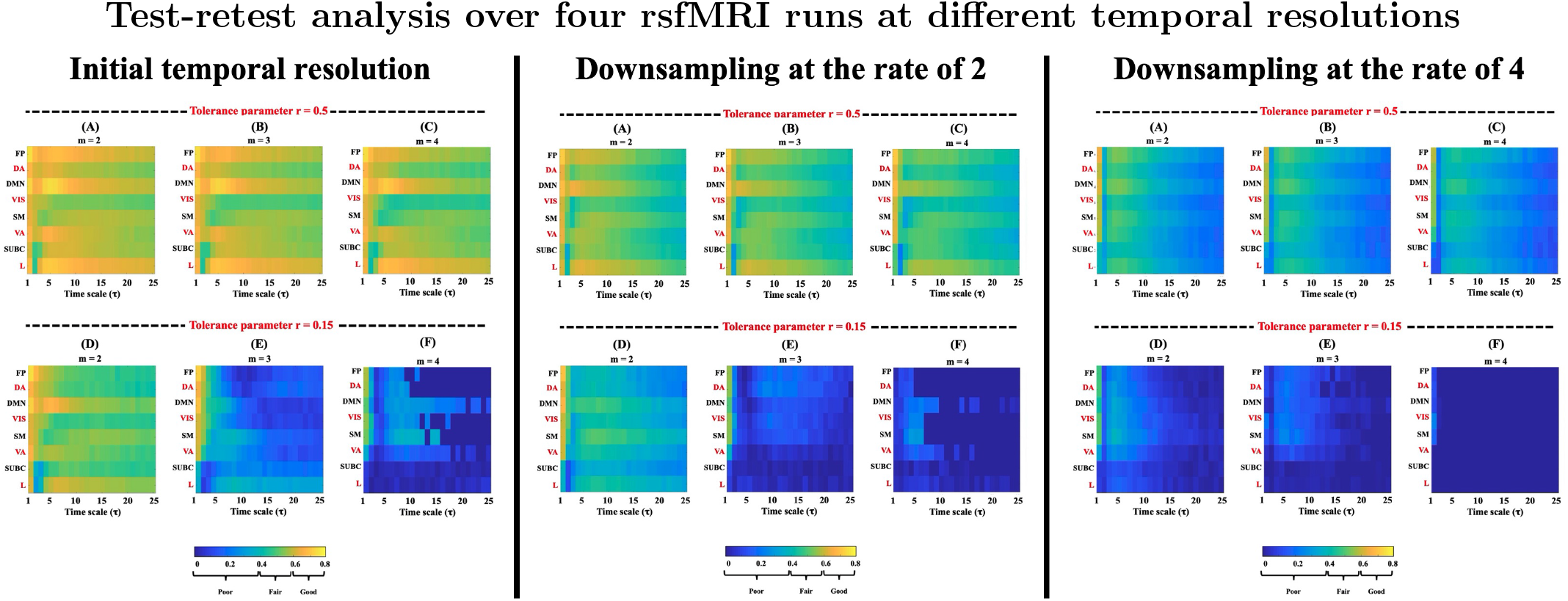
Test-retest analysis of the multi-scale entropy curves of the HCP database over 1000 subjects and four scanning sessions at the embedding dimension *m*=2, the tolerance parameters *r*=0.15,0.5 and three temporal resolutions of rsfMRI. The colors show the intra-class correlation coefficient values ranging from 0 to 1. The values below 0.4 show poor replicability, values between 0.4 to 0.6 show fair replicability, between 0.6 to 0.8 show good replicability and above 0.8 imply excellent reliability (do not exist in the above maps). **Abbreviation**: FP = Frontoparietal, DA = Dorsal Attention, DMN = Default mode network, VIS = Visual, SM = Sensorimotor, VA = Ventral Attention, SUBC = Sub-cortical, L = Limbic. See [51] for the illustrations of RSNs

**Figure S6:**
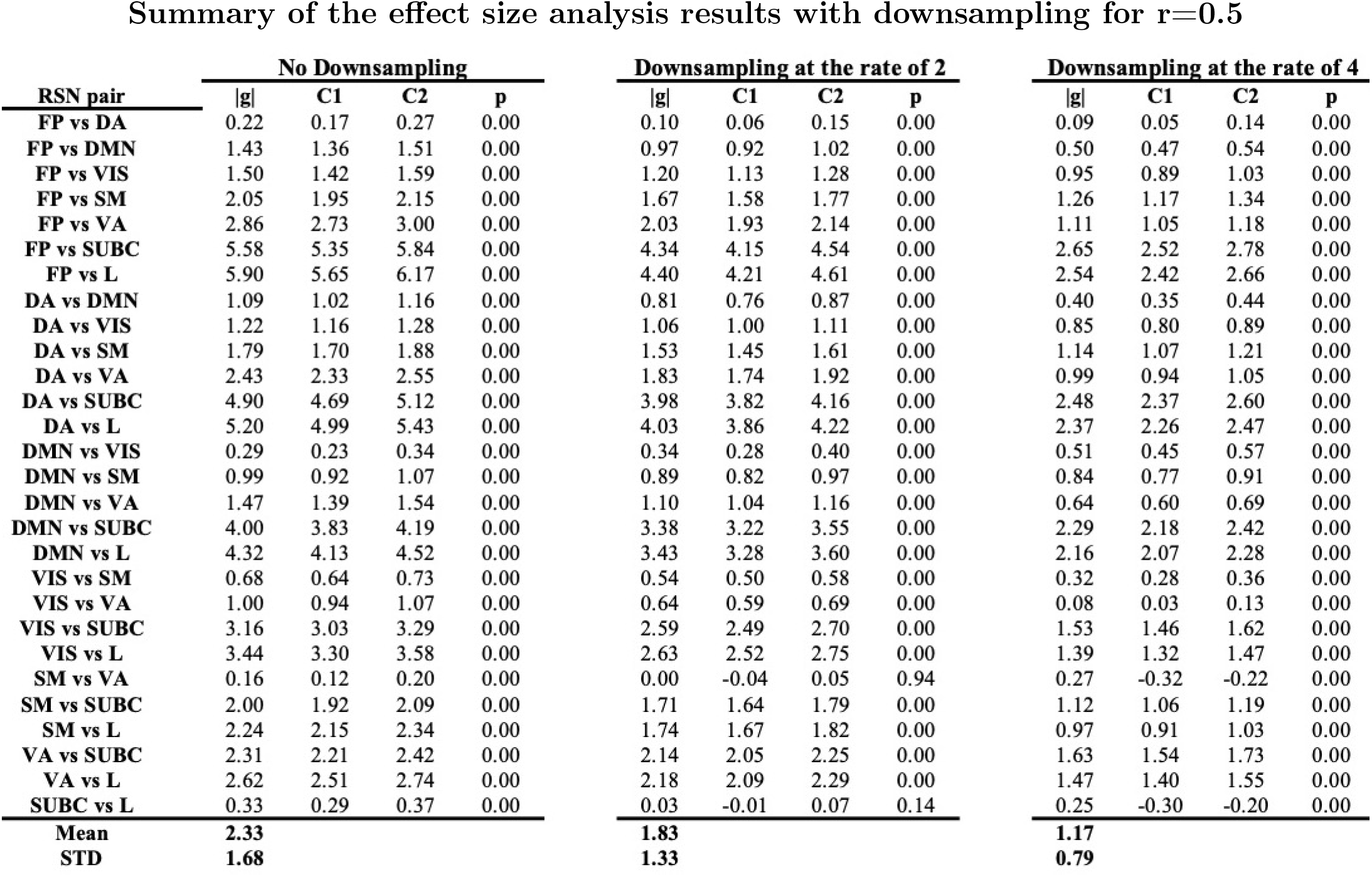
Summary of the effect size analysis of multi-scale entropy results at *r*=**0.5 and** *m*=**2**. In the table, *g* is the Hedges’ *g* measure, *C*1 and *C*2 denote the lower and upper limits of the confidence interval of the Hedges’ *g* after 10000 permutations and *p* represent the associated *p*-value where a value of 0.00 means *p* ≤ 0.001 and corrected for multiple comparisons. **Abbreviation**: FP = Frontoparietal, DA = Dorsal Attention, DMN = Default mode network, VIS = Visual, SM = Sensorimotor, VA = Ventral Attention, SUBC = Sub-cortical, L = Limbic. See [51] for the illustrations of RSNs.

**Figure S7:**
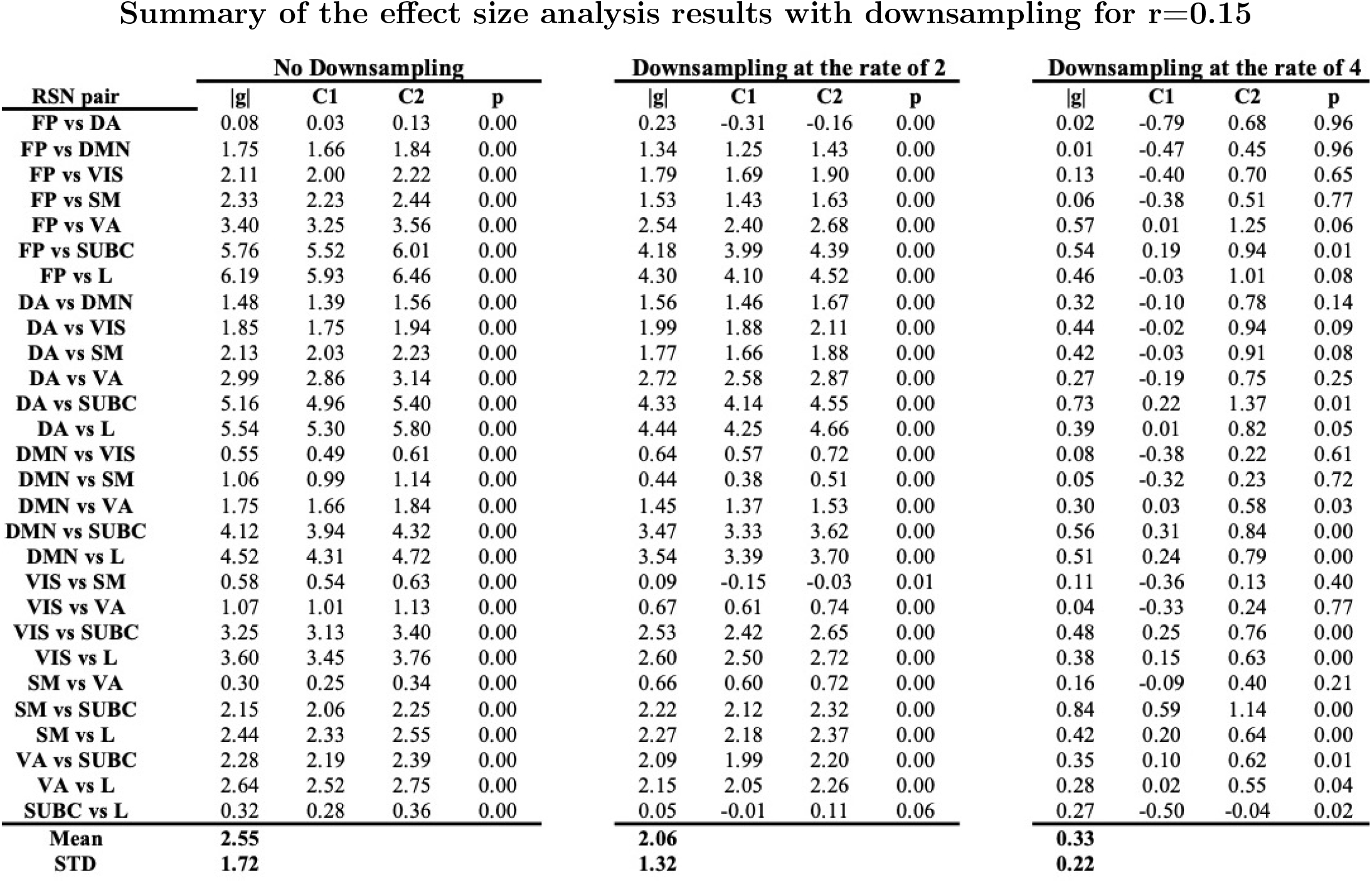
Summary of the effect size analysis of multi-scale entropy results at *r*=**0.15 and** *m*=**2**. In the table, *g* is the Hedges’ *g* measure, *C*1 and *C*2 denote the lower and upper limits of the confidence interval of the Hedges’ *g* after 10000 permutations and *p* represent the associated *p*-value where a value of 0.00 means *p* ≤ 0.001 and corrected for multiple comparisons. **Abbreviation**: FP = Frontoparietal, DA = Dorsal Attention, DMN = Default mode network, VIS = Visual, SM = Sensorimotor, VA = Ventral Attention, SUBC = Sub-cortical, L = Limbic. See [51] for the illustrations of RSNs.

**Figure S8:**
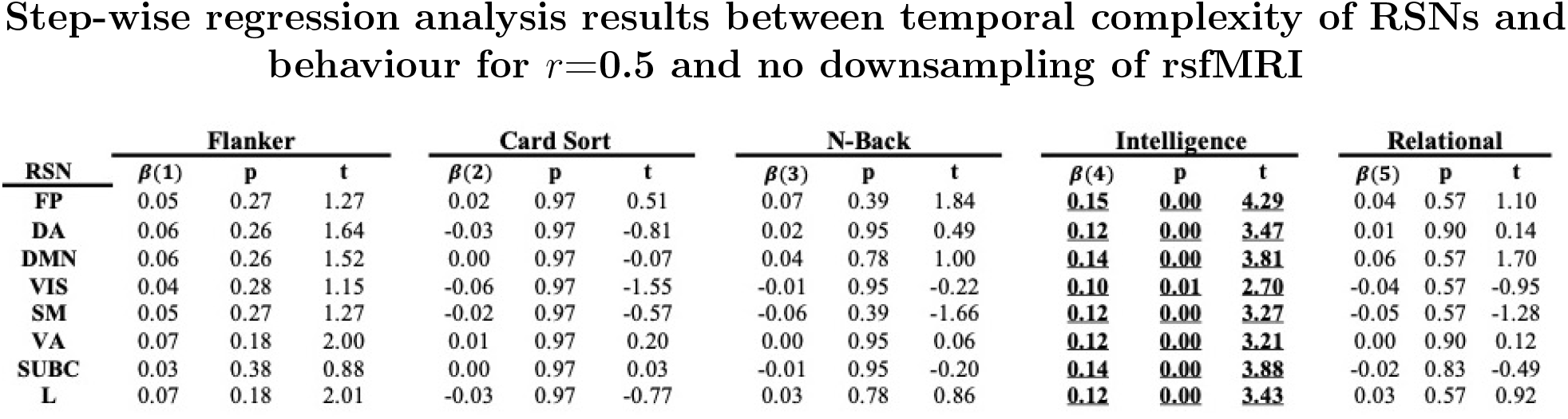
Summary of the multiple regression analysis results between rsfMRI complexity and five HCP behavioural variables for *r*=**0.5**, *m*=**2 and no downsampling** of rsfMRI. The variables are as follows: Variable 1 (*Flanker_Unadj* or the Flanker inhibition measure), Variable 2 (*CardSort_Unadj* or Card Sorting flexibility measure), Variable 3 (*WM_Task_Acc* or N-back working memory measure), Variable 4 (*PMAT24_A_CR* or Ravens fluid intelligence measure) and Variable 5 (*Relational_Task_Acc* or the relational task) [56]. All *p*-values were corrected for multiple comparisons using the false discovery rate method at the significance level of 0.05. Significant *p*-values and their associated *β* coefficients have been highlighted with bold font and underscore. A *p*-value of 0.00 means *p* ≤ 0.001 and corrected for multiple comparisons. **Abbreviation**: FP = Frontoparietal, DA = Dorsal Attention, DMN = Default mode network, VIS = Visual, SM = Sensorimotor, VA = Ventral Attention, SUBC = Sub-cortical, L = Limbic. See [51] for the illustrations of RSNs.

**Figure S9:**
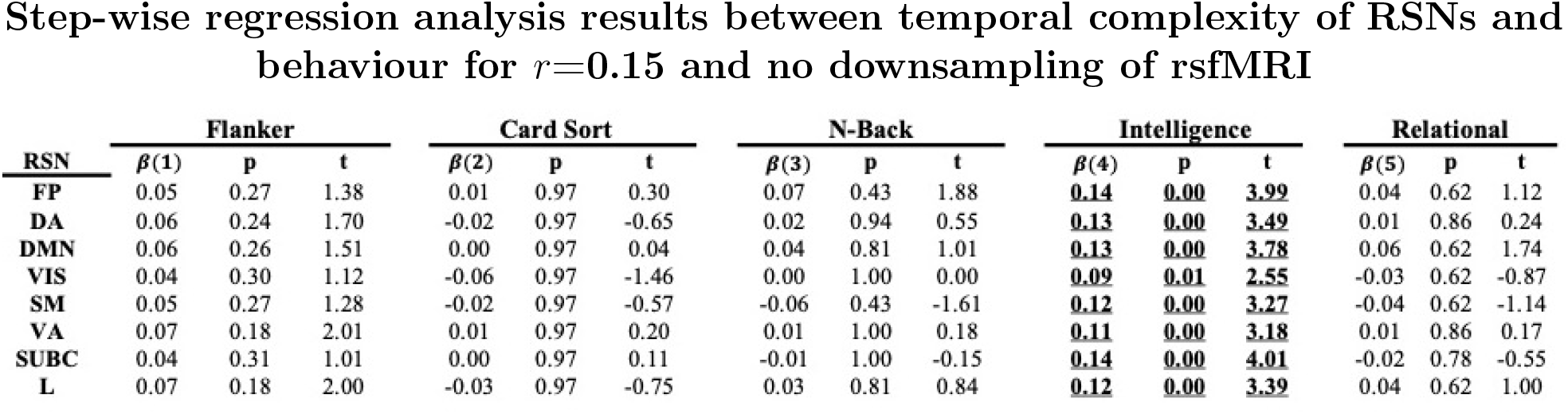
Summary of the multiple regression analysis results between rsfMRI complexity and five HCP behavioural variables for *r*=**0.15**, *m*=**2 and no downsampling** of rsfMRI. The variables are as follows: Variable 1 (*Flanker_Unadj* or the Flanker inhibition measure), Variable 2 (*CardSort_Unadj* or Card Sorting flexibility measure), Variable 3 (*WM_Task_Acc* or N-back working memory measure), Variable 4 (*PMAT24_A_CR* or Ravens fluid intelligence measure) and Variable 5 (*Relational_Task_Acc* or the relational task) [56]. All *p*-values were corrected for multiple comparisons using the false discovery rate method at the significance level of 0.05. Significant *p*-values and their associated *β* coefficients have been highlighted with bold font and underscore. A *p*-value of 0.00 means *p* ≤ 0.001 and corrected for multiple comparisons. **Abbreviation**: FP = Frontoparietal, DA = Dorsal Attention, DMN = Default mode network, VIS = Visual, SM = Sensorimotor, VA = Ventral Attention, SUBC = Sub-cortical, L = Limbic. See [51] for the illustrations of RSNs.

**Figure S10:**
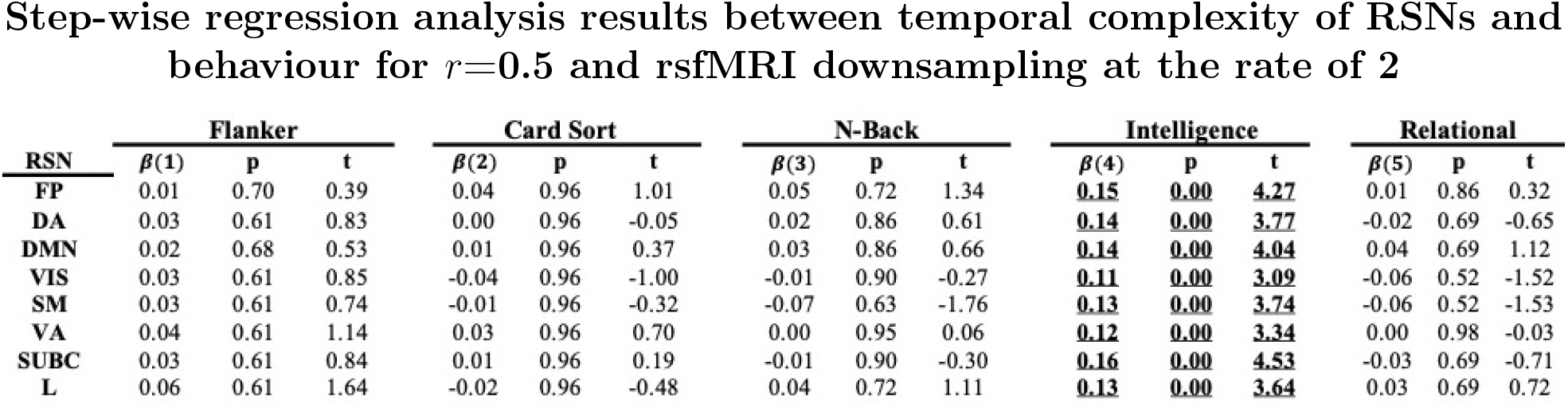
Summary of the multiple regression analysis results between rsfMRI complexity and five HCP behavioural variables for *r*=**0.5**, *m*=**2 and downsampling of rsfMRI at the rate of 2**. The variables are as follows: Variable 1 (*Flanker_Unadj* or the Flanker inhibition measure), Variable 2 (*CardSort_Unadj* or Card Sorting flexibility measure), Variable 3 (*WM_Task_Acc* or N-back working memory measure), Variable 4 (*PMAT24_A_CR* or Ravens fluid intelligence measure) and Variable 5 (*Relational_Task_Acc* or the relational task) [56]. All *p*-values were corrected for multiple comparisons using the false discovery rate method at the significance level of 0.05. Significant *p*-values and their associated *β* coefficients have been highlighted with bold font and underscore. A *p*-value of 0.00 means *p* ≤ 0.001 and corrected for multiple comparisons. **Abbreviation**: FP = Frontoparietal, DA = Dorsal Attention, DMN = Default mode network, VIS = Visual, SM = Sensorimotor, VA = Ventral Attention, SUBC = Sub-cortical, L = Limbic. See [51] for the illustrations of RSNs.

**Figure S11:**
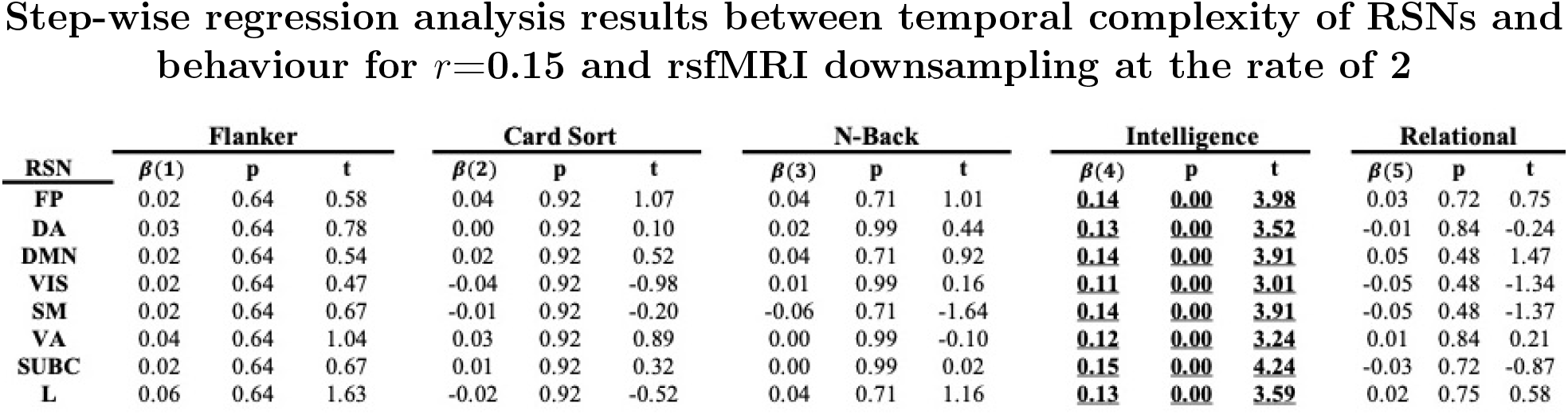
Summary of the multiple regression analysis results between rsfMRI complexity and five HCP behavioural variables for *r*=**0.15**, *m*=**2 and downsampling of rsfMRI at the rate of 2**. The variables are as follows: Variable 1 (*Flanker_Unadj* or the Flanker inhibition measure), Variable 2 (*CardSort_Unadj* or Card Sorting flexibility measure), Variable 3 (*WM_Task_Acc* or N-back working memory measure), Variable 4 (*PMAT24_A_CR* or Ravens fluid intelligence measure) and Variable 5 (*Relational_Task_Acc* or the relational task) [56]. All *p*-values were corrected for multiple comparisons using the false discovery rate method at the significance level of 0.05. Significant *p*-values and their associated *β* coefficients have been highlighted with bold font and underscore. A *p*-value of 0.00 means *p* ≤ 0.001 and corrected for multiple comparisons. **Abbreviation**: FP = Frontoparietal, DA = Dorsal Attention, DMN = Default mode network, VIS = Visual, SM = Sensorimotor, VA = Ventral Attention, SUBC = Sub-cortical, L = Limbic. See [51] for the illustrations of RSNs.

**Figure S12:**
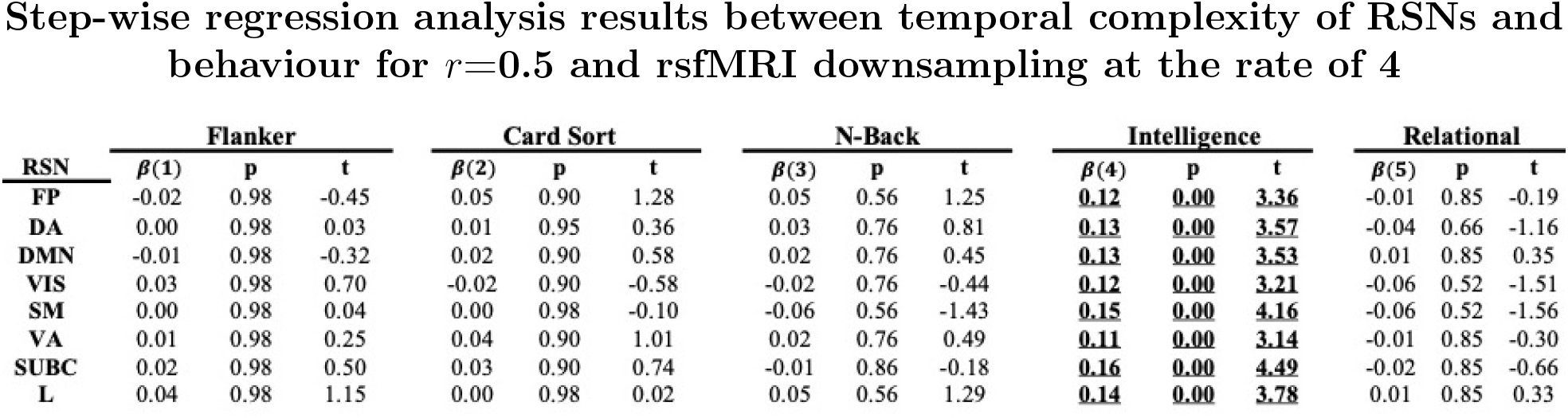
Summary of the multiple regression analysis results between rsfMRI complexity and five HCP behavioural variables for *r*=**0.5**, *m*=**2 and downsampling of rsfMRI at the rate of 4**. The variables are as follows: Variable 1 (*Flanker_Unadj* or the Flanker inhibition measure), Variable 2 (*CardSort_Unadj* or Card Sorting flexibility measure), Variable 3 (*WM_Task_Acc* or N-back working memory measure), Variable 4 (*PMAT24_A_CR* or Ravens fluid intelligence measure) and Variable 5 (*Relational_Task_Acc* or the relational task) [56]. All *p*-values were corrected for multiple comparisons using the false discovery rate method at the significance level of 0.05. Significant *p*-values and their associated *β* coefficients have been highlighted with bold font and underscore. A *p*-value of 0.00 means *p* ≤ 0.001 and corrected for multiple comparisons. **Abbreviation**: FP = Frontoparietal, DA = Dorsal Attention, DMN = Default mode network, VIS = Visual, SM = Sensorimotor, VA = Ventral Attention, SUBC = Sub-cortical, L = Limbic. See [51] for the illustrations of RSNs.

**Figure S13:**
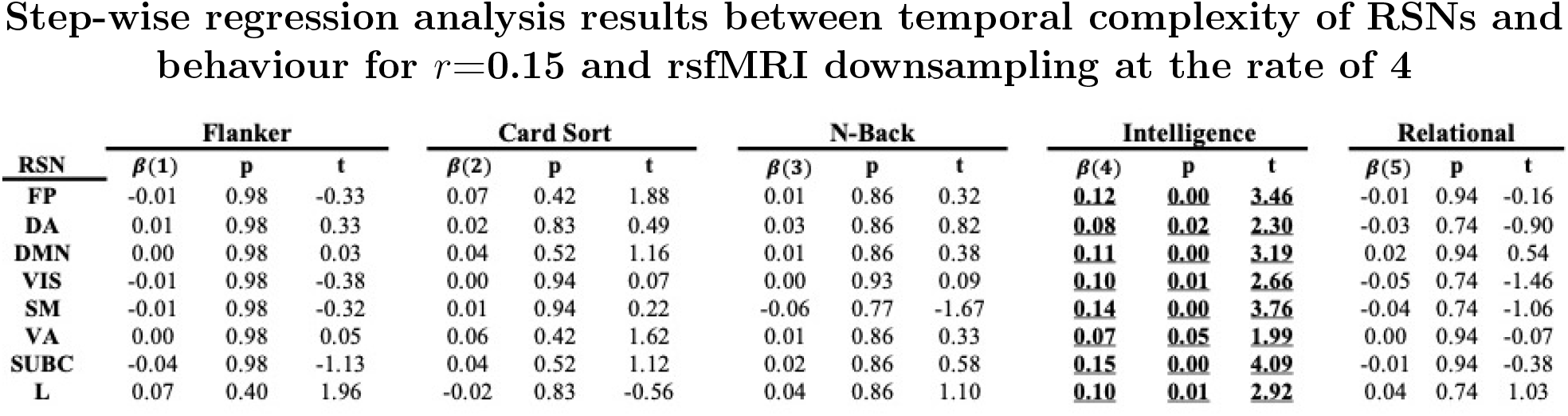
Summary of the multiple regression analysis results between rsfMRI complexity and five HCP behavioural variables for *r*=**0.15**, *m*=**2 and downsampling of rsfMRI at the rate of 4**. The variables are as follows: Variable 1 (*Flanker_Unadj* or the Flanker inhibition measure), Variable 2 (*CardSort_Unadj* or Card Sorting flexibility measure), Variable 3 (*WM_Task_Acc* or N-back working memory measure), Variable 4 (*PMAT24_A_CR* or Ravens fluid intelligence measure) and Variable 5 (*Relational_Task_Acc* or the relational task) [56]. All *p*-values were corrected for multiple comparisons using the false discovery rate method at the significance level of 0.05. Significant *p*-values and their associated *β* coefficients have been highlighted with bold font and underscore. A *p*-value of 0.00 means *p* ≤ 0.001 and corrected for multiple comparisons. **Abbreviation**: FP = Frontoparietal, DA = Dorsal Attention, DMN = Default mode network, VIS = Visual, SM = Sensorimotor, VA = Ventral Attention, SUBC = Sub-cortical, L = Limbic. See [51] for the illustrations of RSNs.

## Footnotes

1 In all equations, scalar variables are in normal font, while vector variables are in bold.

2 The effect size analysis toolbox associated with [53] is available at [the MATLAB File Exchange website](https://au.mathworks.com/matlabcentral/fileexchange/32398-hhentschke-measures-of-effect-size-toolbox).

